# Unsupervised Learning of Brain State Dynamics during Emotion Imagery using High-Density EEG

**DOI:** 10.1101/2020.10.29.361394

**Authors:** Sheng-Hsiou Hsu, Yayu Lin, Julie Onton, Tzyy-Ping Jung, Scott Makeig

## Abstract

Here we assume that emotional states correspond to functional dynamic states of brain and body, and attempt to characterize the appearance of these states in high-density scalp electroencephalographic (EEG) recordings acquired from 31 participants during 1-2 hour sessions, each including fifteen 3-5 min periods of self-induced emotion imagination using the method of guided imagery. EEG offers an objective and high-resolution measurement of whatever portion of cortical electrical dynamics is resolvable from scalp recordings. Despite preliminary progress in EEG-based emotion decoding using supervised machine learning methods, few studies have applied data-driven, unsupervised decomposition approaches to investigate the underlying EEG dynamics by characterizing brain temporal dynamics during emotional experience. This study applies an unsupervised approach – adaptive mixture independent component analysis (adaptive mixture ICA, AMICA) that learns a set of ICA models each accounted for portions of a given multi-channel EEG recording. We demonstrate that 20-model AMICA decomposition can identify distinct EEG patterns or dynamic states active during each of the fifteen emotion-imagery periods. The transition in EEG patterns revealed the time-courses of brain-state dynamics during emotional imagery. These time-courses varied across emotions: “grief” and “happiness” showed more abrupt transitions while “contentment” was nearly indistinguishable from the preceding rest period. The spatial distributions of independent components (ICs) of the AMICA models showed higher similarity within-subject across emotions than within-emotion across subjects. No significant differences in IC distributions were found between positive and negative emotions. However, significant changes in IC distributions during emotional imagery compared to rest were identified in brain areas such as the left prefrontal cortex, the posterior cingulate cortex, the motor cortex, and the visual cortex. The study demonstrates the feasibility of AMICA in modeling high-density and nonstationary EEG and its utility in providing data-driven insights into brain state dynamics during self-paced emotional experiences, which have been difficult to measure. This approach can advance our understanding of highly dynamical emotional processes and improve the performance of EEG-based emotion decoding for affective computing and human-computer interaction.

## 1. Introduction

Emotions are functional brain states internally generated by the brain that give rise to diverse subjective experiences and behaviors in our daily lives (Scherer, 2005). Advancing our understanding of emotions and devising technology for affective computing will enable vast applications in education, healthcare, gaming, and human-computer interaction (Picard, 2000). Despite enormous progress in cognitive science, psychology, and computer science studies of emotions, much research in emotion relies on subjective scores of emotions reported by participants, e.g. verbal expression or questionnaires (Cowen and Keltner, 2017). However, these subjective scores reflect the conscious awareness of their emotional experiences or the expressed emotion, which might differ from the underlying emotional states that could be unconscious or shaped by an individual’s cognitive construct (Picard, 2000; Lindquist et al., 2012; Barrett et al., 2019). To complement the results from subjective reports, objective measurements are used, including behavioral expression (e.g. facial expression (Ekman, 1993)), physiological signals (e.g. electrodermal activity (Sequeira et al., 2009)), and neural signals (e.g. functional magnetic resonance imaging (Kober et al., 2008; Lindquist et al., 2012; Horikawa et al., 2020) and electroencephalography (EEG) (Coan and Allen, 2004; Lin et al., 2010; Kim et al., 2013)). Among these techniques, EEG provides a direct measurement of system-wide brain activities with high temporal resolution, which offers a unique solution to investigating the temporal dynamics of emotional states.

Previous EEG-based emotion studies mostly focused on developing supervised learning approaches to optimize classification accuracy using subjective ratings of emotions as labels, e.g. affective dimensions or emotion categories. Such approaches contributed to the building of emotion classifiers applicable in affective computing and to the identification of EEG biomarkers that maximally separate different labeled emotional conditions. However, these studies often overlooked the temporal dynamics of the emotional states within seconds to minutes of emotional elicitation such as watching videos (Nie et al., 2011), listening to music Lin et al. (2010) or imagining emotional scenarios. For example, Onton and Makeig (2009) conducted a self-paced emotion-imagery experiment where the subjects induced a sequence of 15 distinct emotional states through verbally-guided narratives followed by their own emotional imagery, each lasted 3-5 minutes. A conventional supervised learning approach applied to the dataset assumed the emotional activities were stationary within each 3-5 minutes segment and struggled to achieve high classification accuracy of emotion and to explain the huge difference in classification performance across emotions (Kothe et al., 2013).

Furthermore, most EEG-based emotion studies often assumed that subjective ratings of emotional experiences (e.g. valence and arousal used in popular DEAP dataset (Koelstra et al., 2011)) or objective labels (e.g. from a pool of subjective ratings (Lin et al., 2010)) of emotional stimuli represent the actual emotional states of the individuals. Few studies aimed to validate or challenge these assumptions and explore the relationship between the underlying EEG dynamics and the emotion labels, providing evidence to some of the key questions in the field, such as (a) would the subjective report of emotional experiences be consistent with the objective measurement from neurophysiological recordings, (b) could the difference between emotions be better described by affective dimensions or emotional categories (Barrett, 1998; Mauss and Robinson, 2009; Cowen and Keltner, 2017), (c) what are the temporal dynamics during emotional experiences, and (d) are there specific brain regions associated with specific emotions (Lindquist et al., 2012)

These questions motivate the application of unsupervised learning approaches to characterize emotional states, which attempts to cluster emotional states by EEG activities rather than subjective labels. To our current knowledge, only a few studies have used an unsupervised-learning approach for EEG-based emotion studies. For example, the Gaussian mixture model (GMM) was used for learning segment-level variability in non-stationary EEG Zhuang et al. (2014), a hypergraph representation was introduced to capture the hidden structure of EEG signals across emotion trials (Liang et al., 2019), and the deep belief network (DBN) and hidden Markov model (HMM) were employed for unsupervised feature learning and tracking emotional stage switching (Zheng et al., 2014). However, these studies did not explore the relationship between the hidden structure of EEG and emotion labels.

Alternatively, adaptive mixture independent component analysis (adaptive mixture ICA, AMICA) (Palmer et al., 2008; Hsu et al., 2018a) assumes multidimensional data can be modeled by a mixture of independent component analysis models (ICAMM) (Lee et al., 2000), where each model represents a decomposition of segments of the data into statistically independent components. Previous studies have shown that ICAMM or AMICA can separate EEG activities during different sleep stages (Salazar et al., 2010; Hsu et al., 2018a), fluctuation of drowsiness (Jung et al., 2000; Hsu et al., 2018a), mental states changes during memory test (Safont et al., 2017), guided meditation (Hsu et al., 2018b), and emotional video watching (Ran et al., in press).

This study aims to employ multi-model AMICA as an unsupervised-learning approach for exploring the brain-state dynamics during different emotional experiences. We applied AMICA to a dataset from Onton and Makeig (Onton and Makeig, 2009), containing high-density EEG data (250-channel) collected from 31 subjects during a self-paced emotion-imagery experiment, where the subjects induced a series of 15 emotional states through verbally guided narratives and their own imagination. Marrying a promising unsupervised method for modeling EEG nonstationarity to a unique high-density EEG dataset with a variety of minutes-long emotional experiences enables us to investigate (1) whether such data-driven segmentation is organized and clustered by affective dimensions (e.g. valence) or it is distinct across categories of emotion, and (2) what are the temporal dynamics of EEG during self-paced imagined emotional experience and how do they vary across emotions and subjects. We also reported the plausible neurophysiological sources that activated during emotional imagery and discussed the difference in the AMICA models across emotions and subjects. These results provide evidence that emotional processes could be highly dynamical over time, variable across emotions, and individualized for each subject.

## 2. Materials and Methods

### 2.1. Dataset and preprocessing

#### 2.1.1. Dataset

The dataset used in the study contains 31 recordings collected by Onton and Makeig (2009) and is available at HeadIT website (http://headit.ucsd.edu, *Imagined Emotion with Continuous Data* under *Studies*). EEG data were collected from 250 scalp electrodes with conductive gel using a Biosemi ActiveTwo system (Amsterdam, Netherlands) at a sampling rate of 256 Hz with 24-bit resolution. Caps with a custom whole-head montage were used to position the electrodes, which covered most of the scalp, forehead, and lateral face surface, omitting chin and fleshy cheek areas. Locations of the electrodes for each subject were recorded (Polhemus, Inc.). For more details, see (Onton and Makeig, 2009) and the description on the HeadIT website.

#### 2.1.2. Experiment and Subjects

Thirty-one subjects participated in the emotion-imagery experiment (19 females; age 25.5 ± 5 years) and gave informed consent in accordance with UCSD institutional review board requirements. All subjects reported being able to induce realistic emotional states for most emotions through a verbally guided narrative and their own imagination. Throughout the experiment, subjects were seated comfortably with eyes closed in a dimly-lit room with ear-bud earphones.

Fig. 1(a) shows the paradigm of the self-paced emotion-imagery experiment. Each session started with 2 min of eyes-closed rest in silence, followed by a pre-recorded verbal instructions and a 5-min guided relaxation period to promote a relaxed, inwardly-focused state. In the main task, each subject was presented with a series of 15 recorded voice-guided imagery narratives and was instructed to recall or imagine scenarios for stimulating vivid and embodied experience of the suggested emotion. Subjects were told to take as much time as they needed to recall or imagine a scenario that would induce a realistic experience of the suggested emotion. No external time indicators were provided, and the subjects took on average 3-5 min for each emotional imagery trial. The 15 emotional imagery trials were presented in a pseudo-random sequence alternating between eight positive-valence emotions (love, joy, happiness, relief, compassion, contentedness, excitement, awe) and seven negative-valence emotions (anger, jealousy, disgust, frustration, fear, sadness, grief). Subjects were instructed to press a right-hand button when the emotion began to surge; when the feeling of the emotion subsided, subjects pressed a left-hand button to exit that emotion. Between emotion trials, there was a 1-min rest period consisting of a 40-sec audio clip to guide the subjects’ return to a relaxed, neutral state, followed by a 10-sec silence and then a 10-sec audio clip to help them prepare for the next emotion-imagery trial. The experimental session lasted around 80 min with the duration of each trial varying from 43 sec to 12 min across subjects. The detailed transcripts of the audio clips can be found in the Supplement Information in (Onton and Makeig, 2009).

**Figure 1:**
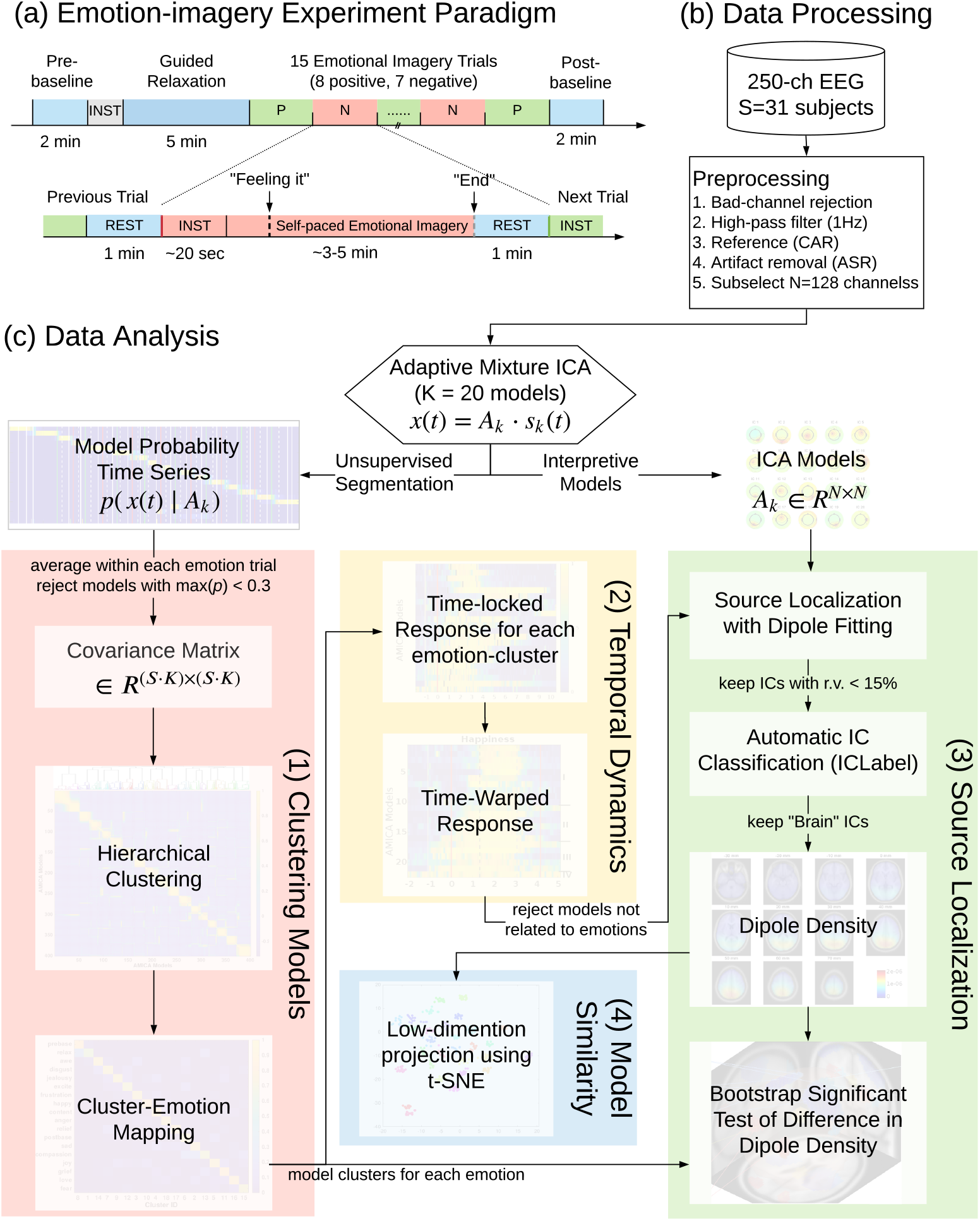
(a) Experimental paradigm of the self-paced emotion-imagery experiment. (b) Flowchart of the EEG data processing prior to the application of the adaptive mixture ICA (AMICA). (c) AMICA model: EEG data, *x*(*t*), are modeled as mixing matrices, *A_k_*, times independent components (ICs) activities, *s_k_* (*t*), for model *k*. Schematic diagram of the post-AMICA data analysis consists of *(1) Clustering Models*: hierarchical clustering of AMICA models across subjects for examining the relationship between emotional-imagery trials and the EEG dynamics segmented by multiple AMICA models, *(2) Temporal Dynamics*: exploring temporal dynamics of emotional responses through time-locked and time-warped analysis, *(3) Source Localization*: mapping ICs from clusters of AMICA models to dipole density for examining differences in source locations during emotional imagery, and *(4) Model Similarity*: projecting the dipole density of individual AMICA models to low-dimension representations through t-distributed stochastic neighbor embedding (t-SNE) for exploring the difference across emotions and subjects.

#### 2.1.3. Data Preprocessing

The data obtained from the HeadIT portal were already pre-processed as described in (Onton and Makeig, 2009) including removal of bad channels (e.g. electrodes with poor skin contact, judged by their grossly abnormal activity patterns), leaving 134-235 channels per subject (214 ± 18, mean ± SD), re-referencing to digitally linked mastoids, and digitally filtered above 1 Hz. Data periods containing broadly distributed, high-amplitude muscle noise and other irregular artifacts were identified by tests for high kurtosis or low probability activity and removed from analysis using EEGLAB functions (Delorme and Makeig, 2004; Delorme et al., 2007). The occurrence of eye blinks, other eye movements, or tonic muscle tension artifacts were not criteria for data rejection.

Further preprocessing steps were applied to the data prior to the application of AM-ICA as shown in Fig. 1(b). Data before the first eye-closed rest session and after the last eye-closed rest session were removed. Data were re-referenced to common average reference. Artifact subspace reconstruction (ASR) (Kothe and Jung, 2016; Mullen et al., 2015) with a cutoff parameter 20 were applied for automatic removal of large-amplitude artifacts like electrode pops and motion artifacts, which has been shown to improve the subsequent ICA decomposition (Chang et al., 2019).

Data were further down-sized to 128 channels by sub-selecting channels that were maximally spaced out in their total spatial distance using *loc-subsets()* function in EEGLAB. The reason was to solve the following three practical issues: insufficient samples for learning a larger number of model parameters (proportional to the square of the number of channels); the computational time was constrained by the resources available; variability of numbers of channels across subjects. The effects of the number of channels on the learning performance of AMICA were further discussed in Section 4.4.

### 2.2. Adaptive Mixture Independent Component Analysis (AMICA)

AMICA assumes that the multi-channel time-series data can be described by a mixture of independent component analysis models. Each model decomposes parts of the data into independent components (IC) and source activities, and each IC has a probability density function parameterized as a mixture of generalized Gaussians. In this study, 20 models, 128 ICs (same as the number of channels), and one generalized Gaussian were used. The choice of 20 models for AMICA was based on an implicit assumption of the underlying states during the emotional imagery experiment, i.e. at least 15 emotions, three baselines, and inter-trial resting periods. The effect of the number of models on the learning performance of AMICA was further addressed in Section 4.1 and has been investigated in Hsu et al. (2018a).

A sphering transformation option was turned on when applying AMICA (*do_pca* =1). Data samples with low probabilities of model fit were rejected (*numreg* = 5, *rejstart* = 2, *rejint* = 5) from being used for learning AMICA parameters to alleviate the effects of transient artifacts such as electrode pops and discontinuities. AMICA employs the expectation-maximization (EM) algorithm to estimate the parameters that maximize the data likelihood, using an efficient implementation with parallel computing capability (Palmer et al., 2008) available at https://github.com/japalmer29/amica and as an open-source plug-in for EEGLAB (Delorme and Makeig, 2004). The computation jobs were run on the Comet Supercomputer at San Diego Supercomputer with the support of the Neuroscience Gateway (NSG) (Sivagnanam et al., 2013). The max learning iteration was set to 2000 steps, which took around 48 hours on one computing node with 24 threads. A retrospective analysis showed the data likelihood reached a plateau around 1000 steps. For more details on the model, learning algorithm, effect and selection of learning parameters, please see Section S1.1 in the Supplemental Materials and Palmer et al. (2008) and Hsu et al. (2018a).

It is worth noting that AMICA failed to learn and returned a data likelihood of zero when applying to 2 out of the 31 recordings (Subject IDs 26 and 32) regardless of the number of models and channels used. This might be due to the numerical round-off errors when computing a data likelihood close to zero. Since the re-implementation of AMICA on NSG was not available, these two recordings were removed from further analysis.

As an unsupervised approach with the assumption of the ICA mixture model, AM-ICA learns the underlying data clusters and can quantitatively assess continuous changes in the EEG patterns reflecting the state transitions. Specifically, the activation of each ICA model can be represented as the normalized data likelihood given the estimated parameters of each model, referred to as “ICA model probability” that indicates the goodness-of-fit of the ICA model to each data sample. Furthermore, AMICA also provides interpretable models allowing the characterization of the spatial distribution and frequency content of active sources in each brain state.

### 2.3. Post-AMICA data analysis

#### 2.3.1. Clustering AMICA models across subjects

To characterize the relationship between AMICA models and emotion states, we clustered the AMICA models with similar activation patterns across subjects. The procedure is summarized in Fig. 1(c) *(1) Clustering Models* and detailed below. An 18-by-1 feature vector was created for each model which consisted of its average model probability from emotion surge button presses to the end of the emotion trial for each of the 18 states (15 emotion states plus guided relaxation, pre-session baseline, and post-session baseline). The models with low averaged probability (i.e. below 0.3) in any of the 18 states were rejected for further analysis, resulting in 401 out of 580 (20 models × 29 subjects) models remaining. The correlations (ρ) between each pair of these feature vectors were computed, yielding a correlation matrix of the size 401× 401. The agglomerative hierarchical clustering with 18 clusters was applied to the correlation distance matrix (1 – *ρ*) to cluster models with highly-correlated feature vectors across subjects, using *linkage()* with group average and *dendrogram()* functions in MATLAB. The feature vectors within each model cluster were averaged to obtain the final cluster-emotion mapping graph for examining the relationship between the emotional-imagery trials and the EEG dynamics segmented by multiple AMICA models.

#### 2.3.2. Time-locked and time-warped temporal dynamics of model transition

Fig. 1(c) (2) *Temporal Dynamics* illustrated the procedure to explore the EEG temporal dynamics during emotional imagery. The model probability time-series of the AMICA models in the cluster identified according to section 2.3.1 were smoothed with 5-second non-overlapping windows, time-locked to the start of the emotion-imagery trial from 2 minutes before to 8 minutes after this event. To address the variability in trial lengths across subjects, the model-probability time courses were linearly time-warped to the median reaction time (i.e. start to button press) and median trial lengths (i.e. start to end of the trial) such that these events were aligned across subjects, using the *timewarp()* function in EEGLAB (Delorme and Makeig, 2004).

#### 2.3.3. Criteria for categorizing model types based on activation time courses

To help identify and interpret emotion-associated models based on the temporal dynamics of their probability time courses, we categorized models into four types, as illustrated in Fig. 3(b). For each model, we calculated the average model probabilities in the four time periods during the emotion-imagery trial: *P_pre_*: from 2-min before to the end of the previous trial; *P_rest_*: from the end of the previous trial to the start of the current trial; *P_instruct_*: from the start of the current trial to the first button press; *P_feel_*: from the button press to the end of the current trial.

The specific criteria for the four model types are: Type IV: *P_pre_* > 0.5, “activated” (increased probability) since the previous emotion-imagery trial; Type III: excluding Type IV, the remaining models with *P_rest_* > 0.5, meaning the model activated during the rest period and before the start of the trial; Type II, the remaining models with *P_instruct_* > 0.5, indicating the model activated after the start of the trial when the guided audio clip began; and Type I, the remaining models with *P_feel_* > 0.5 - those only activated after the button press through the end of the trial.

These model types provide a semi-quantitative way to compare temporal dynamics across emotions. The distribution of model types for each emotion-associated model cluster was summarized in Table 1. For the subsequent source localization and dipole density analyses, the models with types III and IV were removed because they activated before the start of the trials, indicating the responses might not be associated with the specific emotional-imagery trial.

**Table 1:** Types of models for each of the fifteen model clusters with their corresponding emotions. The criteria for categorizing models are described in Section 2.3.3

| Emotion \ Types | I | II | III | IV | Total | I+II (%) | III (%) | IV (%) |
| --- | --- | --- | --- | --- | --- | --- | --- | --- |
| grief | 10 | 8 | 0 | 2 | 20 | 90 | 0 | 10 |
| excitement | 9 | 8 | 3 | 2 | 22 | 77 | 14 | 9 |
| happiness | 10 | 6 | 5 | 1 | 22 | 73 | 23 | 5 |
| love | 4 | 9 | 5 | 0 | 18 | 72 | 28 | 0 |
| fear | 10 | 7 | 5 | 2 | 24 | 71 | 21 | 8 |
| relief | 6 | 7 | 4 | 2 | 19 | 68 | 21 | 11 |
| anger | 8 | 9 | 3 | 5 | 25 | 68 | 12 | 20 |
| sadness | 7 | 5 | 4 | 2 | 18 | 67 | 22 | 11 |
| disgust | 8 | 4 | 4 | 2 | 18 | 67 | 22 | 11 |
| awe | 9 | 6 | 6 | 2 | 23 | 65 | 26 | 9 |
| joy | 7 | 3 | 5 | 2 | 17 | 59 | 29 | 12 |
| compassion | 2 | 5 | 2 | 3 | 12 | 58 | 17 | 25 |
| jealousy | 4 | 6 | 5 | 3 | 18 | 56 | 28 | 17 |
| frustration | 3 | 7 | 7 | 4 | 21 | 48 | 33 | 19 |
| contentment | 2 | 4 | 8 | 8 | 22 | 27 | 36 | 36 |
| average | 6.6 | 6.3 | 4.4 | 2.7 | 19.9 | 64 | 22 | 13 |

#### 2.3.4. Source localization with dipole fitting

To characterize and compare the spatial distribution of ICs for each AMICA model in the model cluster, we applied source localization using DIPFIT2 (Acar and Makeig, 2010), available in EEGLAB. The electrode locations were manually co-registered to the template head model (4-layer Boundary Element Model). The single equivalent current dipole in the head model was fit to each IC given its spatial projection to the scalp electrodes. Coarse and fine grid search for the best-fitting dipole locations were applied in sequence using DIPTFIT2. Non-dipolar ICs with residual variance larger than 15% or those ICs with dipole locations outside of the head model by a 5-mm margin were removed from further analysis. On average, 60% of the ICs were retained across subjects.

#### 2.3.5. Automatic IC classification

To investigate and interpret the AMICA models, we categorized the independent components (ICs) of all the models into seven types, namely Brain, Eye, Muscle, Heart, Channel Noise, Line Noise, and Others, using an automatic IC classifier (ICLabel) (Pion-Tonachini et al., 2019), which is a pre-trained neural network based on the power spectra and spatial distribution of the ICs. To improve the performance of ICLabel for multimodel AMICA, we weighted the power spectra of ICs by the activation (i.e. the normalized log-likelihood) of the model to which the ICs belonged. The Lite version of ICLabel was used which was faster and yielded comparable results with that of the default version. The code and detailed tutorial can be found in https://github.com/sccn/ICLabel.

#### 2.3.6. Dipole density computation and bootstrap significance testing

The dipole density of the selected ICs (i.e. with r.v.<15%, inside of the head model, and classified as “Brain” ICs) from all models in the same cluster were obtained by summing over each IC’s contribution using a spatial Gaussian-kernel with full width at half maximum (FWHM) of 8.5-mm, weighted by its total power, and normalizing to 1. The *dipoleDensity()* function in EEGLAB was used. The total power of each IC was computed with 4-sec non-overlapping windows, weighted by the average model probability in the same time window of the model to which the IC belonged. The dipole density was computed for each AMICA model for the t-SNE visualization described in Section 2.3.7.

Furthermore, we used bootstrapping to test the significant differences between dipole density pairs of different emotion model clusters. Two surrogate datasets were generated by drawing two sets of ICs randomly with replacement from the pool of ICs from all models in the two clusters. The number of ICs in each surrogate dataset equaled the size of each original emotion dataset. The dipole densities of the two surrogate datasets were computed (described in Methods 2.3.4), and their difference was obtained. After repeating this process 1000 times, the distribution of surrogate differences for each voxel provides a difference value above which only 5% of the surrogate values fell, indicating a significance of 0.05 if the original dataset difference falls above that value.

To characterize brain sources activated during emotional imagery, we also combined all models activated during the emotional-imagery trials in one group (Emotion group) and compared to all models activated during relaxation, pre-session baseline, and post-session baseline (Baseline group). The same bootstrapping approach was applied to obtain the significant difference in dipole densities between the Emotion and Baseline groups, with 1000 bootstrap repetitions.

For better visualization, the statistically significant dipole density was plotted in a reconstructed 3D image using code snippets from the Measure Projection Toolbox (MPT) (Bigdely-Shamlo et al., 2013). Each 2×2×2 mm voxel was color-coded by the dipole density difference ranging from −1000 to 1300 *IC/mm*^3^. The figure also shows the 2D projections on the sagittal, coronal and axial MR images for a more comprehensive view. The areas of significance were labeled with their Broadmann areas (BA) defined in MNI coordinates using the MNI to Talairach mapping application in the Yale BioImage Suite Package https://bioimagesuiteweb.github.io/webapp/.

#### 2.3.7. Visualization of model similarities using t-distributed stochastic neighbor embedding (t-SNE)

The t-distributed stochastic neighborhood embedding (t-SNE) method (van der Maaten and Hinton, 2008) was used to visualize the similarity between the dipole densities of each AMICA model by projecting the 91 × 109 × 91-dimensional dipole density down to two-dimensional space. The t-SNE is a nonlinear dimensionality-reduction method that finds a map in low-dimensional space reflecting pair-wise similarities of neighboring points in the high-dimensional space by minimizing the Kullback-Leibler divergence between the two distributions in the two spaces. The *tsne()* function from MATLAB was used with the Euclidean distance metric.

## 3. Results

### 3.1. Unsupervised segmentation of state changes during emotional imagery

Applied to high-density EEG, the 20-model AMICA was able to characterize and separately model the EEG dynamics underlying brain-state changes during the emotion-imagery experiment. Fig. 2a shows the model probabilities, i.e. normalized data log-likelihood given to each of the 20 models, over experimental time for a sample subject. Model 1 (M1) was the only model with high model probabilities during the initial instruction and pre-session baseline periods. Around the time the “relaxation” audio clip was played, M1 faded and M2 emerged and became the best-fitted model. Starting with the first emotion-imagery trial, M3 became the dominant model after the “happiness” audio clip was played (the solid green line) and continued until the end of the trial (the solid white line) and before M4 became the dominant model in the next “fear” trial. This transition of dominant models sometimes happened around the start of the audio clips (solid green lines for positive emotions and red lines for negative emotions), but for other emotion trials such as “excitement”, “contentment”, “anger”, and “grief”, the transitions were time-locked to the button presses indicating the target emotion was felt (dashed white lines). In this case, AMICA could separately model the EEG activity of almost all emotional imagery trials (except the “disgust” trial) and resting baselines, meaning a nearly one-to-one mapping between AMICA models and the emotional imagery trials.

**Figure 2:**
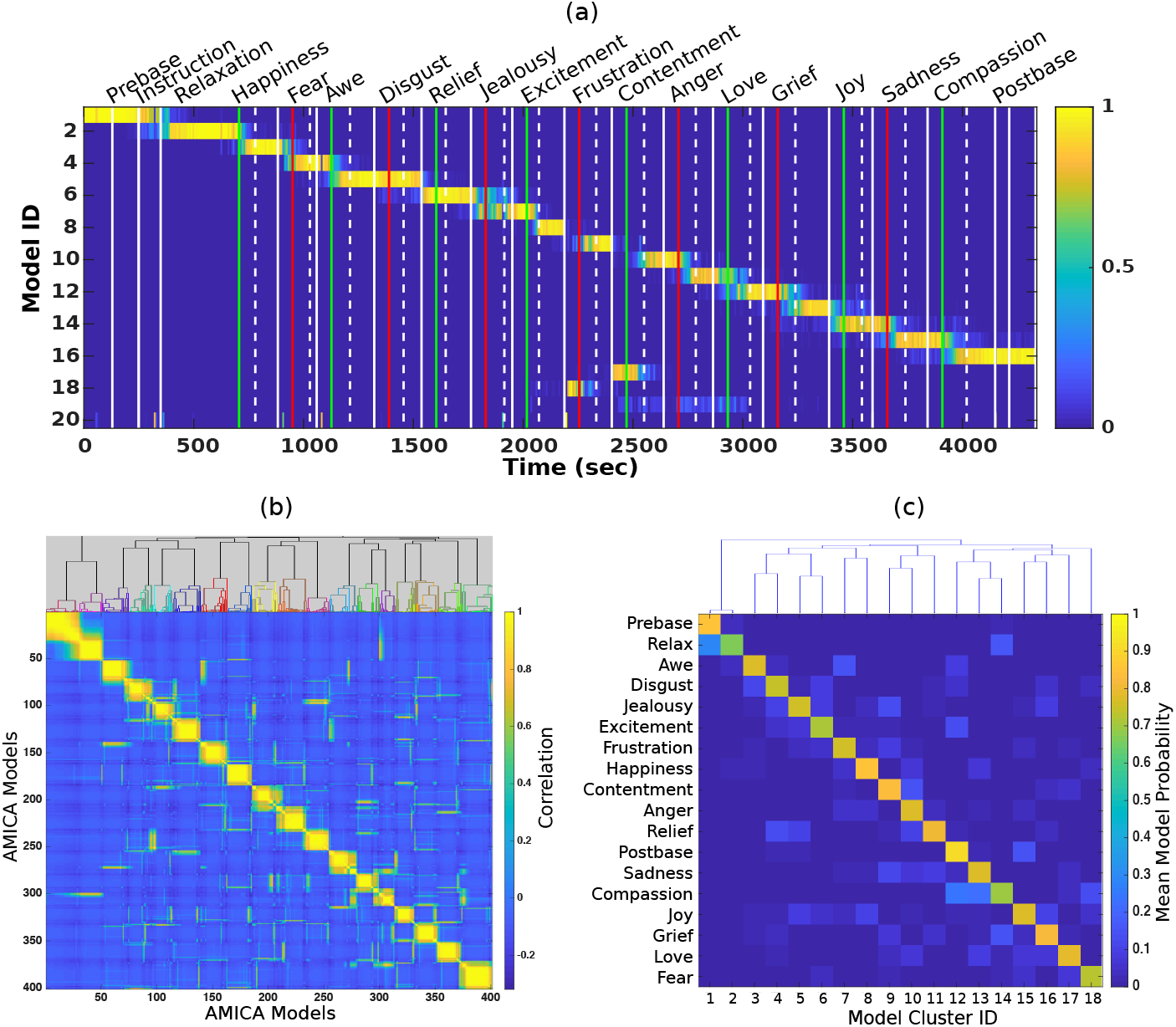
(a) Probability time series of 20 models learned by AMICA for a single subject. The models were sorted based on their active time in the experiment. The vertical color lines indicate the start of each emotional imagery trial with positive (green) and negative (red) emotions. The dashed and solid white lines indicate the button presses when the subject felt the emotion and when the subject exited the trial, respectively. (b) Pair-wise correlation coefficients between all AMICA models from all subjects using model activation patterns, i.e. averaged probability within each of the 18 segments (15 emotion trials plus relaxation and two baseline periods), sorted according to the hierarchical clustering result with 18 clusters. The dendrogram of the clustering result is shown at the top where different colors represent each model cluster. (c) The average model activation patterns for each model cluster, with the dendrogram representing distances between the clusters shown at the top.

The mapping between AMICA models and emotional imagery trial markers was observed across most of the subjects. The probability time series of 20-model AMICA from all subjects were included in Figs. S1 to S5 in the Supplemental Materials. Across subjects, the 20-model AMICA segmented EEG activity into 10 to 18 states with an average of 14 states, each with at least one minute of activation, i.e. averaged probability larger than 0.5 in a 5-sec smoothing window. The rest of the models were only active sporadically or during short periods of resting between two emotional imagery trials.

### 3.2. EEG activity in different emotional imagery trials could be modeled by distinct AMICA models

Hierarchical Clustering (HC) was applied to identify models with similar emotion-related activation patterns across subjects, e.g. models with high probability in the same emotion-imagery trial(s). Fig. 2b shows the pair-wise correlation between all models across all subjects, sorted by the HC result shown at the top of the panel. Each of the 18 clusters contained 14 to 30 models. The result of a block-diagonal pattern shows a clear separation between model clusters and high similarity within each model cluster, suggesting that models with similar activation patterns could be consistently found across subjects.

Furthermore, the averaged activation patterns found within each model cluster (each column in Fig. 2c) only corresponded to one of the 18 experimental segments. That is, these models in the cluster from different subjects were only activated in one of 15 emotional-imagery trials, two baselines, or relaxation periods. This mapping between model clusters and their corresponding segments was almost one-to-one. A few exceptions were pre-session baseline (“prebase”) and “relax” for Cluster 1 and 2, and “sadness” and “compassion” for Cluster 13 and 14, in which the clusters were closer in distance according to the HC result shown at the top of the figure. Hence, the multi-model AMICA decomposition was able to distinguish the EEG dynamics of each emotional-imagery trials, allowing HC to identify model clusters that corresponded to distinct emotion categories across subjects.

### 3.3. Temporal dynamics of model transitions during emotional imagery varied across subjects and emotions

Once the best-fitted models for each emotion were identified across subjects, we examined their model-probability time series as a means to characterize the temporal dynamics of state transition during emotional imagery. Fig. 3a shows the 5-sec smoothed probability time series of the AMICA models in Cluster #8 (Fig 2c) which were active during the “Happiness” emotion trials, time-locked to the beginning of the trial - the onset of the audio clip (red line). The subjects’ reaction time, defined as the duration from the onset of the audio clip to subjects’ button presses (black line), varied from 0.6 to 7.2 minutes across subjects; while the total duration of the trials, from the onset of the audio clip to the exit event (gray line), varied from 1.5 to 9.4 minutes.

**Figure 3:**
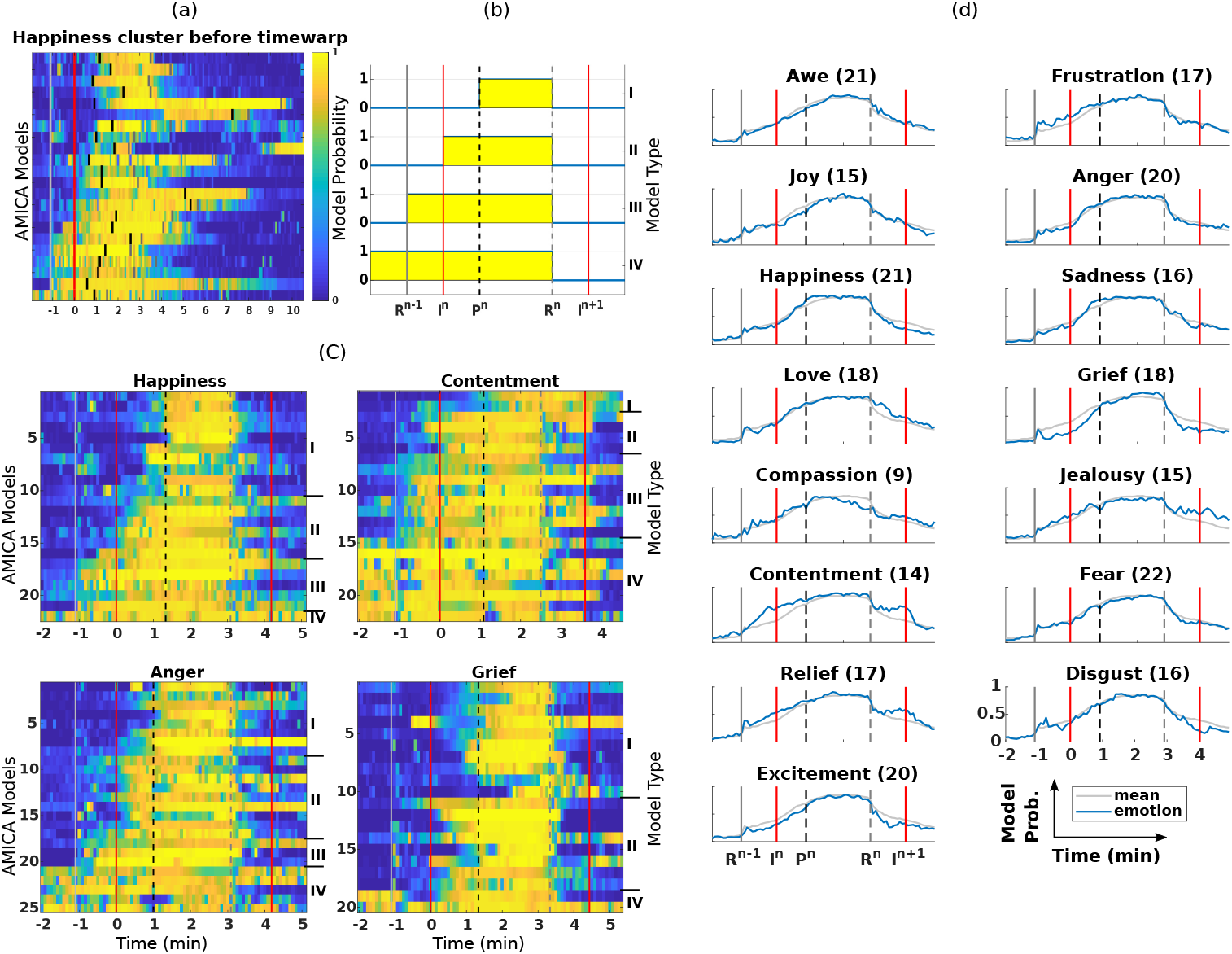
(a) Model probability time series of all AMICA models across subjects in the same cluster activated during imagined happiness trials, time-locked to the beginning of the trials. Vertical lines mark the end of the previous emotion trial (white, *R*^n-1^), the onset of the audio instruction (red, *I*^n^), the first button press when the subject felt the emotion (black, *P*^n^), and the end of the current trial (gray, *R^n^*). (b) Illustration of model types I to IV based on the timing of their activation patterns. Detailed criteria were defined in Section 2.3.3. (c) The model-probability time-series plots for happiness, contentment, anger, and grief trials. The “feeling it” button press (dashed black line) and the end of the current emotion trial (dash gray line) were time-warped to the median press latencies from all subjects whose model contributed to the cluster. The models were sorted according to their types. See Table 1 for detailed information. Results from all 15 emotions were shown in Fig. S6 and S7 in the Supplemental Materials. (d) The median model-probability time series across subjects for each emotion cluster (blue curve) and the average time series across all emotions (gray curve), time-locked to the start of the resting period (gray vertical line) and the beginning of the emotion trial (red). The button presses and the end of the emotion trial were time-warped to the median values across subjects and marked as dashed black and gray lines respectively. The number of models for each cluster was indicated in the parentheses. Models related to positive (left column) and negative emotions (right column) were sorted by the median values of the trial length.

To better visualize transitions between active AMICA models during emotional imagery trials, we time-warped the probabilities of the AMICA models between button presses (dashed black line) and the exit events (dash gray line) to their median values across subjects, as shown in Fig. 3c for “happiness”, “contentment”, “anger”, and “grief”. Results from all 15 emotions were shown in Fig. S6 and S7 in the Supplemental Materials. We found that the models became active (transition from low (blue) to high (yellow) model probability) at different times during the emotion-imagery trials for different subjects. Based on this observation, the models were further categorized into four types based on their activation patterns shown in Fig. 3b. Some model transitions were time-locked to the button presses (Type I) or the onset of the audio clip (Type II). Other models were active before the onset of the audio clip, as soon as the previous emotion trial ended (Type III), while other models even had a high probability starting from the previous emotion trials (Type IV). The criteria for categorizing the four types of models are described in Section 2.3.3.

Fig. 3c shows that the activation patterns as described by model types varied significantly across emotions. For instance, 45% and 50% of the models in the “happiness” and “grief” clusters, respectively, were Type I, which were the highest percentages among all the emotion clusters. This indicates the EEG activities during these emotions were distinct from the previous emotion and the rest period and were not induced quickly after listening to the audio clip. The “Anger” cluster had more Type II than Type I models, suggesting the response might happen as soon as the subjects heard the audio clip. Type III and Type IV models were the most common in “contentment” (36% were Type III and Type IV, respectively), suggesting the EEG activities during imagined contentment were similar to those during the resting period or even the previous emotion trials.

The numbers and percentages of the four types of models for each of the 15 emotions are summarized in Table 1. Across all emotions, on average 64% of models were Type I and II, which were more likely related to the imagined emotions given that their occurrence time-locked to the start of the audio instruction or the button press. The percentages ranged from 27% for “contentment” to higher than 70% for “grief” and “happiness”. On average, 22% of models were Type III, suggesting the EEG activities learned by the models during the emotional imagery trials resembles those during the preceding rest period. Lastly, 13% of models were Type IV, which were unlikely to be related to the emotion-specific activity, given that they were also active during the previous emotion - either the subjects continued to feel the previous emotion or the new emotion was not successfully elicited. Therefore, types III and IV models were rejected from further source-localization analysis.

Fig. 3d summarizes the temporal dynamics of state transitions for each emotion, showing the median values of model probability across all models in the same emotion cluster for each of the 5-sec windows during the emotion-imagery trials. The grand averages of the model probabilities time series of all emotions were shown as gray curves. Type IV models were excluded given that they did not likely capture the emotion-related activities.

For all emotions, similar inverted U-shaped probability time courses were found, which indicate the beginning, middle, and end of the emotional imagery trials. The onset, slope and duration of the rising phase, the plateau, and the falling phase of the probability time courses varied across emotions. Small increases in the model probability started as early as the beginning of the resting period after the previous emotion trial ended (gray line). The rising phase continued, and for some emotions the slope increased after the beginning of the audio instruction (red line). The time courses nearly reached the peaks when the subjects felt the emotion and pressed the button (dashed black line) and then plateaued until they pressed the button again indicating they were ready to end the trial (dash gray line). The model probability immediately decreased after subjects exited the trial and entered the rest period. Cycles of these temporal dynamics lasted on average for 4-6 minutes.

Comparing the temporal dynamics across emotions, we found that the rising phases of the probability time series between *I*^n^ and *P*^n^ had higher slopes for emotions like happiness, sadness, anger, and grief, compared to the average response (gray curve). The probability time series before *P*^n^ and after *R*^n^ were lower than average for emotions like grief and excitement and were higher for emotions such as contentment and relief. For happiness, excitement, and most of the negative emotions, the model probabilities dropped faster than average after the end of the emotion trials (*R*^n^); while emotions such as love and contentment tended to last longer.

### 3.4. AMICA models learned high percentage of dipolar brain ICs

Among all the models in the emotion clusters, Table 2 shows that on average 24.4±8.6% of the 128 ICs from each model were classified as Brain ICs, while only 6.5±3.8% and 3.1 ±2.8% of ICs were classified as muscle-related and eye-related ICs, respectively, using an automatic IC classifier - ICLabel (Pion-Tonachini et al., 2019). The remaining ICs were mostly Others (44.6±8.5%).

**Table 2:** (Top) Mean and standard deviation of percentage of ICs in each category classified by ICLabel across all AMICA models. (Bottom) Percent of dipolar ICs (with r.v.<15%) within each class and among all dipolar ICs.

| Percentage (%) | Brain | Muscle | Eye | Heart | Line Noise | Channel Noise | Other |
| --- | --- | --- | --- | --- | --- | --- | --- |
| Mean Percent | 24.4 | 6.5 | 3.1 | 0.3 | 13.8 | 7.3 | 44.6 |
| STD | 8.6 | 3.8 | 2.8 | 0.4 | 5.2 | 4.4 | 8.5 |
| Dipolar ICs | 92.2 | 31.5 | 43.0 | 62.7 | 74.1 | 29.5 | 48.1 |
| Total dipolar ICs account for | 37.3 | 3.5 | 2.5 | 0.4 | 16.8 | 3.7 | 35.8 |

To validate the quality of IC decomposition by multi-model AMICA, we examined the percentage of the dipolar ICs of each model (i.e. matching the projection of a single equivalent dipole less than a given residual variance), as a quality metric suggested by Delorme et al. (2012). We found that on average 60% of the ICs were dipolar, defined as less than 15% of residual variance from their counterparts generated by a single dipole model. Among the dipolar ICs, 37% of ICS were labeled as Brain ICs, while only 3.5% and 2.5% of ICs were labeled as Muscle and Eye ICs.

We also found that the AMICA models rejected from the clustering analysis had a lower percentage of Brain ICs (20%) and a higher percentage of Other ICs (49%). However, there was no significant difference in the percentage of Brain ICs between models from different emotion clusters.

### 3.5. Dipole densities of the AMICA models

To quantitatively compare the brain sources across AMICA models, we computed the dipole density of each AMICA model, excluding non-dipolar ICs (r.v.<15%) and non-Brain ICs (by ICLabel). See Section 2.3.5 for details. Fig. S8 in Supplemental Materials shows the dipole density of dipolar brain ICs from all AMICA models during baseline periods and relaxation. We found that most of the ICs were located at the bilateral occipital and central parietal regions.

The t-distributed stochastic neighborhood embedding (t-SNE) was used to visualize the similarity between dipole densities of AMICA models by projecting the 91×109×91-dimension dipole-density features down to two dimensions. Fig. 4 shows that dipole distributions were far more similar across emotions within the same subject than the same emotion across subjects.

**Figure 4:**
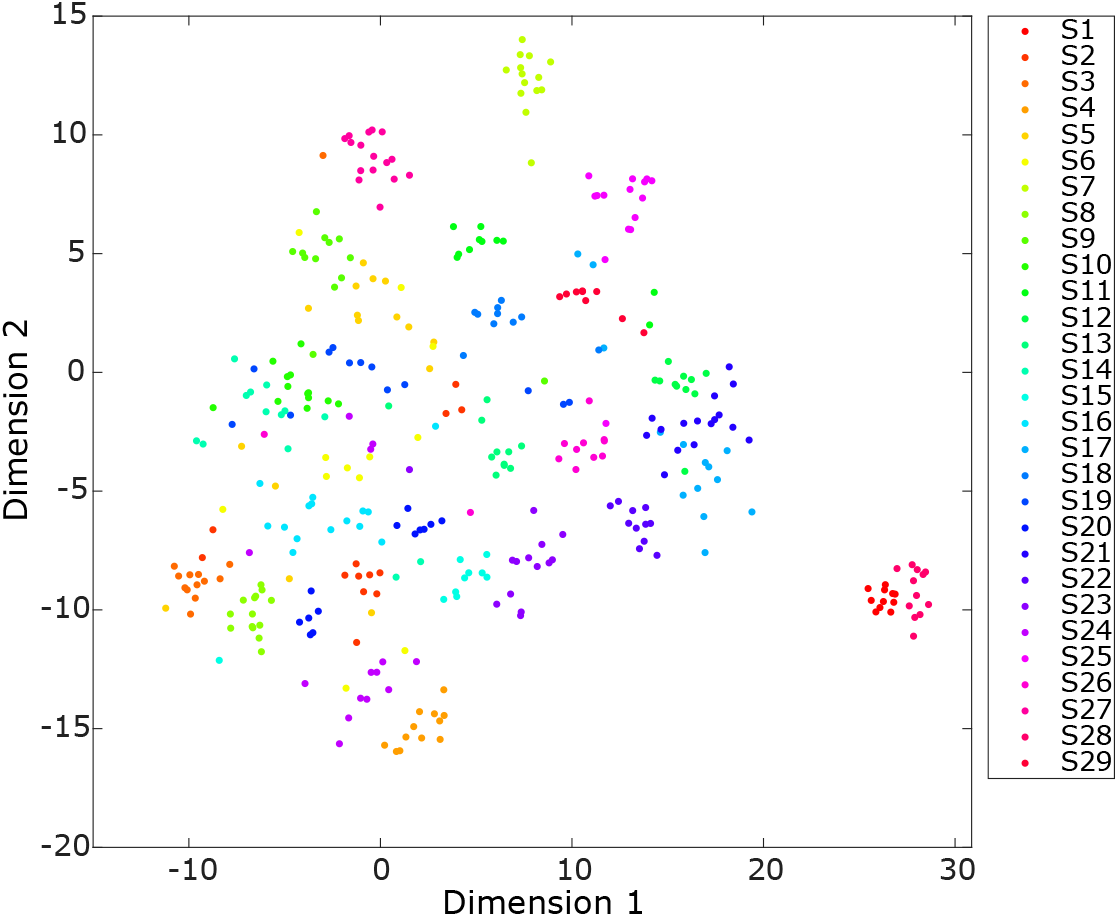
Projecting dipole densities of all AMICA models onto a 2-dimensional feature space using t-SNE, color-coded by subject index.

### 3.6. Spatial distribution of brain sources activated during emotional imagery

When we first compared the dipole density of AMICA models between pair-wise emotion clusters, we found that the difference in dipole density could be biased by which subjects’ models were selected in the clusters, due to the variability of dipole density results across subjects (as shown in Fig. 4) and the limited number of models in each cluster (e.g. “contentment” cluster only had six type I and II models for the dipole density analysis, Table 1). To address the subject-selection bias, we pooled together model clusters from the eight positive emotions (97 models) and compared the aggregated dipole density from that of model clusters from the seven negative emotions (96 models) using bootstrapping (see Section 2.3.6). However, no significant difference in dipole density was found between the positive and negative emotion models.

Finally, we pooled together AMICA models from all 15 emotion clusters (“Emotion cluster”) and compared the aggregated dipole density from that of model clusters from the three baseline periods (“Baseline cluster”), namely the pre- and post-baseline and relaxation. Fig. 5 shows significant reductions in dipole density in the Emotion cluster compared to the Baseline cluster in the premotor cortex (Brodmann Area (BA) 6), primary somatosensory cortex (BA 1), and primary motor cortex (BA 4), with the right hemisphere showing a broader effect than the left hemisphere. Additionally, Fig. 5 shows slight reductions in dipole density in the Emotion cluster in the left dorsolateral prefrontal cortex (BA 9), left anterior prefrontal cortex (BA10), and right insula (BA13). In contrast, the Emotion cluster showed significant increases in dipole density in the associative visual cortex (BA 19), left angular gyrus (BA 39), and ventral posterior cingulate cortex (BA 23).

**Figure 5:**
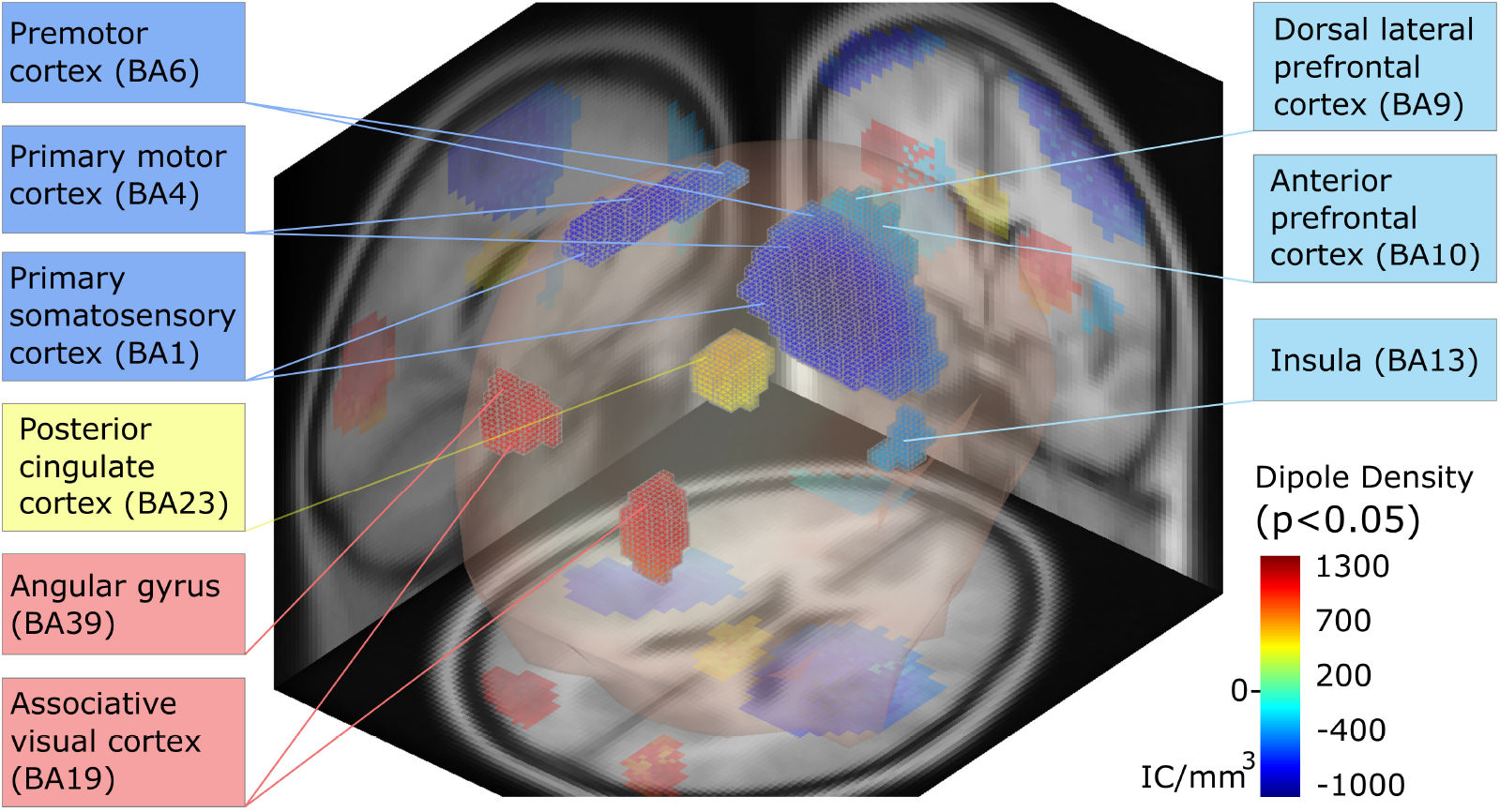
The difference in dipole densities of dipolar brain ICs of the AMICA models accounted for all emotions versus baseline periods (i.e. pre- and post-session baseline and relaxation), masked by the result of a significance test (*p* < 0.05) using bootstrapping. The result is superimposed on a 3D brain model and the sagittal, coronal and axial slices of a template MR image. The corresponding Brodmann areas (BA) were identified by their MNI coordinates.

## 4. Discussion

This study aims to explore the often-overlooked EEG dynamics during emotional experiences. The uniqueness of the study is that we applied multi-model AMICA as an unsupervised approach to high-density EEG data recorded during a self-paced emotional imagery experiment where a variety of minutes-long emotional activities were elicited. This enables us to investigate (1) whether AMICA can identify meaningful clusters of EEG dynamics during emotional imagery and how do such clusters correspond to different emotions (e.g. dimensional or categorical)? (2) what are the temporal dynamics of EEG during the self-paced imagined emotional experience and how do they vary across emotions and subjects? and (3) what are the neurophysiological sources that activated during emotional imagery?

### 4.1. Data-driven AMICA approach separates distinct EEG activities across imagery of 15 emotional scenarios

We first showed that multi-model AMICA was able to reliably resolve 10 to 18 distinct states throughout the experiment across 29 subjects where the EEG activities in each state could be separately modeled by one of the ICA mixture models (Fig. 2a). Furthermore, the unsupervised segmentation of these states clearly corresponded to the onset and offset of each emotion-imagery trial. The results suggest that EEG activities during the imagery of 15 different emotional scenarios (plus two baseline and relaxation periods) were distinct and separable from each other by AMICA without event or label information.

Interestingly, our results did not show EEG patterns clustered according to affective dimensions of emotions such as valence or arousal. Rather, for each of the 18 experimental segments including 15 emotional imagery trials and three baseline periods, AMICA could learn a distinct cluster of models across subjects where all models in the cluster were only active (i.e. highly probable to model the data) in that segment (Figure 2c). Only in a few cases, the identified model clusters were activated in more than one experimental segments (with the second segment has averaged probability larger than 0.3), such as “prebase” and “relax” and also “sadness” and “compassion”.

While our main results show that the EEG activities could be distinctly modeled for each of the 15 emotional categories when 20-model AMICA was applied, one ought to be cautious to conclude that emotion activities were organized categorically. We conducted a pilot analysis to examine whether the activities of some emotions may be more similar compared with that of other emotions by applying AMICA with various numbers of models to the same EEG dataset. The result from a sample subject (Fig. S9) shows that when 5 models were used, several consecutive emotional imagery trials shared the same model (e.g. pre-session baseline, relaxation and awe by Model 1 and compassion, fear, contentment, and jealousy by Model 5). When the number of models increased to 10, we observed that several models were active in two consecutive trials with the same valence (e.g. frustration and anger by Model 2, joy and happiness by Model 3, sadness and grief by Model 4, love, relief and excitement by model 5, etc.). As the number of models increased to 15, the separation between each emotional imagery trial was clearer. When the number of models reached 20, almost every trial was distinctly separated by different models. The combined evidence seemed to suggest that, even though EEG activities of each of the emotional imagery trials could be distinctly separated when a large number of models were available, the trials that had the same valence might share more similar EEG activities that could be modeled together when only a smaller number of models was used. Further investigation is needed to systematically examine this effect across subjects.

### 4.2. Temporal dynamics of emotional activities and variability across emotions

AMICA also enables the characterization of the temporal dynamics of EEG patterns with a sub-second resolution, inferred by the changes of models that could best fit the data. We found that the transition of the best-fitted models aligned well in time with different phases of the emotional imagery trials (Figure 3). Among the models that were dominant in each emotional imagery trial, on average, one-third of the models became active when the subjects listened to the narratives (Type II models) and one-third of the models became active when the subjects pressed the button and indicated “feeling the emotion” (Type I models) (Table 1). Most of the dominant models “faded away” upon the subjects ending the trials and starting the rest period. These dominant models wax and wane with the onset and offset of the emotional imagery trials, providing strong evidence that the models separated EEG activities related to the emotional states.

Comparing the temporal dynamics of different emotional states, we found that emotion trials such as grief, excitement, and happiness presented more distinctive emotional activities, evidenced by a higher percentage of Type I and II models (Table 1 and Figure 3). Some emotional states such as love, contentment, and the relief lasted longer after the end of the imagery trials. These temporal features could be used as new measures in understanding and characterizing differences in emotional activities.

We also found that the last one-third of dominant models became active during the rest period prior to the emotional imagery trial (Type III models) or even during the previous emotion-imagery trial (Type IV models). One explanation is that the emotional activities were not successfully induced because subjects became fatigued over the long experiment or reported failure in producing certain emotions, especially jealousy and compassion which had a higher percentage of Type III and IV models (Table 1). Another reason could be that the induced emotional states were similar to that during the prior rest period such that the brain activity was largely similar as well. For example, “contentment” is likely to be close to the resting state and, indeed, “contentment” had the highest percentage of Type III and IV models (73%).

These objective EEG results suggest that distinctive emotional states might not always have been achieved during the emotional imagery task, providing evidence for the low accuracy in emotion classification obtained in a previous study using the same dataset (Kothe et al., 2013). In the study, Kothe et al. (2013) applied a machine learning model to a subset (12 subjects) of the same EEG dataset with a goal to predict the positive versus negative valence of a novel (e.g. previously unseen) emotion experienced by the same subject. They found that the classification accuracies were the highest for emotional imagery trials of grief and happiness (higher than 90%), while the accuracies were the lowest for the trials of awe, jealousy, contentment, and frustration (lower than 65%) (see Fig. 1 in Kothe et al. (2013)). The results of their classification accuracies for different emotions highly correlated with the percentages of Type I and II models those emotions had in this study (Table 1). In other words, if the emotional states were not successfully elicited or were less distinctive from the resting state, as evidenced by a lower percentage of Type I and II models, these emotional states had poor classification accuracy – close to the chance level. This poses AMICA as a promising tool to assess whether, and when, emotional activities are elicited for studying emotion (e.g. comparing emotion-eliciting paradigms) and improving emotion classification performance (e.g. selecting best time window).

### 4.3. Neurophysiological sources activated during emotional imagery and inter-subject variability

Toward a neurophysiological interpretation of AMICA models, we first examined the quality of the AMICA decomposition and found that on average 60% of ICs were dipolar (i.e. can be fitted with a single equivalent dipole with less than 15% residual variance). The high percentage of dipolar ICs is correlated with high quality of ICA decomposition and is comparable as reported in Delorme et al. (2012). This provides evidence for the validity of ICs from multi-model AMICA decomposition and allows for dipole density analysis. Further application of an automatic IC classifier, ICLabel (Pion-Tonachini et al., 2019), revealed that on average 37% of the dipolar ICs were brain-related independent components (ICs) (Table 2), suggesting AMICA models were able to identify multiple independent brain processes. Dipolar and brain ICs were selected for computing the dipole density plot. A sample dipole density plot of dipolar brain ICs from the Baseline model clusters (Fig. S8) shows that those dipolar brain sources were mostly located in the occipital and parietal cortices, consistent with those observed during eyes-closed resting, as the subjects also closed their eyes in the emotional-imagery experiment.

Visualizing dipole densities of all AMICA models in a 2D t-SNE feature space (Fig. 4) revealed that the dipole distributions were more similar across emotions within the same subject (i.e. recording) compared to those with the same emotion across subjects. This implies the brain from a particular subject might have a core set of ICs that do not change significantly between models within the same AMICA decomposition. It is, however, these minor differences that define the specificity of models for different emotions within-subject.

The minor differences in dipole distributions across emotions were not consistent enough to rise to significance across subjects. Furthermore, when we pooled together model clusters from all of the positive emotions and compared them to those from all of the negative emotions, the bootstrap test did not show any significant difference (*p* > 0.05) in dipole density between the two. This result is consistent with a seminal review paper on neuroimaging evidence of emotion (Lindquist et al., 2012), where their meta-analytic analysis found “little evidence that discrete emotion categories can be consistently and specifically localized to distinct brain regions”.

However, it is important to point out alternative explanations to the non-significant dipole-density result between emotions. First, it could be speculated that emotions might be too individualized to exhibit consistent IC expressions across subjects. Secondly, differences among emotional states may be due to changes in activation time-series of ICs in addition to IC distributions (Onton and Makeig, 2009). AMICA models include parameters for learning the probability density function (PDF) of the activation time-series of each IC. The spectral contents of the IC activation may also explain the differences between models. One of the challenges for comparing PDFs or spectral contents of IC activations across emotions is the identification of common IC clusters across emotions since models from different subsets and numbers of subjects were included in different emotion model clusters. Although this study could not show unique patterns of IC distributions for individual emotions across subjects, future work applying AMICA to a larger number of subjects could potentially overcome the inter-subject variability and identify differences in dipole density between emotions.

Despite the non-significant dipole-density result between emotions and the vast inter-subject differences in the spatial distributions of ICs, we found significant differences in dipole density when experiencing emotions compared to baseline (resting) periods, which were consistent across subjects. These were brain areas that came on- or offline as the brain moved from neutral to active emotional imagery and processing. The primary somatosensory cortex, primary motor cortex, and premotor cortex, which are related to motor and sensory functions, showed significant decreases in dipole density (Figure 5) during emotional imagery. The left dorsolateral prefrontal cortex (dlPFC), left anterior prefrontal cortex (aPFC), and right insula also showed significant decreases in dipole density. Previous studies found that the dlPFC is associated with emotion processing and emotion regulation (Morawetz et al., 2016; Viviani, 2014; Kerestes et al., 2012; Ray and Zald, 2012; Aupperle et al., 2012). The aPFC is believed to be related to memory recall and coordinating information processing (Ramnani and Owen, 2004), and specifically the left aPFC was found to be activated during emotion-regulation tasks (Koch et al., 2018; Bramson et al., 2018; Morawetz et al., 2017). Recent studies found that the anterior part of the insula could be related to emotion recognition and emotional awareness (Motomura et al., 2019; Zhang et al., 2019; Craig, 2009). In addition, significant increases in dipole density during emotional imagery were found at the associative visual cortex, which is related to the processing of visual information, and the left angular gyrus, which is related to the processing of visually perceived words and might be associated with memory retrieval (Seghier, 2013). The other region that showed significantly more IC expressions was the posterior cingulate cortex (PCC), which could be associated with various brain functions including memory retrieval and attentional focus (Rolls, 2019; Leech and Sharp, 2014).

Taken together, these brain areas activated or deactivated during emotional imagery were largely consistent with previous neuroimaging studies. In the review paper, Lindquist et al. (2012) reported a set of brain regions commonly involved during emotion experience across discrete emotion categories. This includes the motor cortex supporting language and executive attention, the visual cortex connecting with areas involved in core affect like the amygdala, and the prefrontal cortex and medial posterior group involved with conceptualization (Kober et al., 2008).

These results, revealed by the analysis of the dipole distributions derived from AMICA models corresponding to specific emotion or baseline periods, may require a new interpretation relative to traditional ICA models. Conventional ICA models use all data points from an entire experiment, which usually include resting periods between trials and between blocks. As such, ICA trained on both resting and active brain states. The single ICA model that is derived is then examined in the power spectrum dynamics to observe changes occurring when the brain transitions from resting to task behavior. The activity that most drives the ICA solution is a high-amplitude synchronous activity, like a resting alpha rhythm. Therefore, often it is possible to observe an ICA component activation changes from high-amplitude alpha activity during rest to low-amplitude gamma activity during task performance, even though low-amplitude gamma is not usually a driving force in ICA decompositions. In contrast, AMICA finds unmixing matrices for specific points in time, rather than the entire recording. In our case, AMICA found solutions that often described single emotion periods. This means that the parts of the cortex that were “active” for a given emotion may be invisible to ICA because the prevailing activity was low-amplitude gamma activity that does not typically drive ICA decompositions. Therefore, when we see a decrease in dipole density in AMICA models during emotion periods compared to baseline, this may very well indicate patches of cortex that became engaged for the task. This interpretation would explain the decrease in dipole density we observed in the dlPFC and aPFC. The increases in dipole density in the PCC and associative visual cortex would be interpreted using the same logic, but taking into account the functions and characteristics of those brain regions. In the visual cortex, the most parsimonious explanation is that active emotional experiencing resulted in larger alpha activity in the visual cortex. This result is somewhat surprising as the visual imagery is known to decrease alpha activity. One explanation may be that while emotional imagery required some visual imagery, the relaxation narrative that played during our baseline period may have resulted in even more vivid visual imagery. In the cingulate cortex, the “active” frequency range is in the theta (4-7 Hz) range, which is a synchronized high-amplitude rhythm, rather than the gamma range. Thus, an increase in dipole density in this region may indicate heightened processing in this region during emotional imagery.

### 4.4. Limitations and future works

Although AMICA successfully modeled EEG activities and characterized brain-state changes during the emotional imagery experiment, it did not prove that the models were generalizable across subjects. It may be that verbally guided imagery of emotion could be confounded with the motor, auditory, visual, or other sensory imagery suggested. To disentangle such effects, future work can include a control condition with only motor or sensory imagery and test whether the emotional imagery trial and the control condition share the same AMICA model.

Since the sub-selected 128-channel EEG montage for training AMICA models covered the whole scalp and partial facial and neck areas (i.e. “whole montage”), it could be argued that the facial and neck muscle artifacts significantly contributed to the separation of EEG activities between different emotion-imagery trials. To test the hypothesis, when we sub-selected a different subset of 128 channels only from electrodes placed above the ears (i.e. “scalp montage”), the AMICA decompositions showed consistent results in terms of separating EEG segments that corresponded to the emotion trials. In addition, the IC classification of the 128-channel whole montage using ICLabel shows that the numbers of artifactual ICs such as eye, heart, and muscle components were much smaller than the number of brain ICs (Table 2). The evidence suggests that the unsupervised segmentation of AMICA was not solely due to the contribution of muscle and eye artifacts, especially from facial and neck muscles.

For a neurophysiological interpretation of the AMICA models, we used dipole density to map the high-dimensional AMICA parameters of each model onto the same brain space. This enables systematic comparison and statistical testing of spatial distributions across AMICA models. However, this mapping involves user-defined thresholds for the selection of dipolar ICs and types of ICs, which would affect the resulting dipole density. Alternative approaches to systematically compare and quantify the distance between the AMICA models in a high-dimensional parameter-space would advance the interpretability of AMICA results and provide further neurophysiological insights.

As discussed in the previous section, AMICA models - albeit able to separate different emotions - seem to be individualized and specific to each subject. This could be the result of over-fitting, given that there were approximately 338K parameters to learn (for 20 AMICA models each with 128 ICs) with 1.38M data samples (for a 90-min recording). In fact, we have tried to further sub-select 64 channels for training AMICA models. The empirical result showed that the AMICA models with 64 ICs could still segment the EEG data in a consistent way, but the transition between segments (i.e. changes in model probability) was not as distinct compared with the results from 128- and 250-IC AMICA.

An important future direction would be testing whether the emotion-associated AMICA models can be transferred to EEG activities of a separate test session, from different subjects, or variable emotion-eliciting paradigms. Although training AMICA models imposes a heavy computational burden, effectively trained models could be used to make statistical inference on test data in a near-real-time fashion, enabling online emotion decoding.

## 5. Conclusions

This study aims to provide a new perspective in studying emotion by data-driven discovery of brain-state dynamics during imagery of emotional experiences. We applied an unsupervised-learning approach, AMICA, to learn a mixture ICA models that elucidate a high-density EEG dataset collected from 31 subjects during a self-paced emotion-imagery experiment. We first showed that EEG data were mostly distinct and separable (without label information) during subjects’ imagery of 15 different emotional experiences. The state transition in EEG dynamics revealed the time-courses of emotional activities which corresponded to subjective report (i.e. button press) of the emotional feeling. The time-courses varied across emotion: for example, “grief” and “happiness” showed more abrupt state transitions during the onset and offset of emotional imagery, “contentment” was almost indistinguishable from the resting period, and “love” lasted longer after the end of imagery. Comparing the results reported in the previous study (Kothe et al., 2013) revealed that emotion with less distinctive state changes in EEG activities during imagery had poor emotion classification accuracy.

These models learned a high percentage of dipolar brain components mostly distributed in the occipital and parietal regions. The spatial distributions of dipolar brain ICs of the AMICA models showed higher similarity within-subject across emotions than within-emotion across subjects. No significant difference in the dipole densities was found between positive and negative emotions. However, significant changes in IC distributions during emotional imagery compared to rest were identified in the left dorsal lateral and anterior prefrontal cortex, posterior cingulate cortex, right insula, motor cortex, and visual cortex. This study demonstrates the use of AMICA as an unsupervised approach in learning the underlying brain state dynamics in an emotion imagery experiment, shedding light on the neurophysiological underpinning of emotional experiences, and thereby improving the performance of emotion decoding for EEG-based affective computing and advancing our understanding of emotion.

## 6. Acknowledgments

This work was supported in part by the U.S. National Science Foundation (IIP-1719130) and the Army Research Laboratory (W911NF-10-2-0022). Dr. Makeig’s participation was funded by a grant from the U.S. National Institutes of Health (R01 NS047293-13A1) and by a gift from The Swartz Foundation (Old Field, NY).

## References

Acar, Z.A., Makeig, S., 2010. Neuroelectromagnetic forward head modeling toolbox. Journal of neuroscience methods 190, 258–270.

Aupperle, R.L., Allard, C.B., Grimes, E.M., Simmons, A.N., Flagan, T., Behrooznia, M., Cissell, S.H., Twamley, E.W., Thorp, S.R., Norman, S.B., et al., 2012. Dorsolateral prefrontal cortex activation during emotional anticipation and neuropsychological performance in posttraumatic stress disorder. Archives of general psychiatry 69, 360–371.

Barrett, L.F., 1998. Discrete emotions or dimensions? the role of valence focus and arousal focus. Cognition & Emotion 12, 579–599.

Barrett, L.F., Adolphs, R., Marsella, S., Martinez, A.M., Pollak, S.D., 2019. Emotional expressions reconsidered: Challenges to inferring emotion from human facial movements. Psychological Science in the Public Interest 20, 1–68.

Bigdely-Shamlo, N., Mullen, T., Kreutz-Delgado, K., Makeig, S., 2013. Measure projection analysis: a probabilistic approach to eeg source comparison and multi-subject inference. Neuroimage 72, 287–303.

Bramson, B., Jensen, O., Toni, I., Roelofs, K., 2018. Cortical oscillatory mechanisms supporting the control of human social–emotional actions. Journal of Neuroscience 38, 5739–5749. doi:10.1523/JNEUROSCI.3382-17.2018.

Chang, C.Y., Hsu, S.H., Pion-Tonachini, L., Jung, T.P., 2019. Evaluation of artifact subspace reconstruction for automatic artifact components removal in multi-channel eeg recordings. IEEE Transactions on Biomedical Engineering.

Coan, J.A., Allen, J.J., 2004. Frontal eeg asymmetry as a moderator and mediator of emotion. Biological psychology 67, 7–50.

Cowen, A.S., Keltner, D., 2017. Self-report captures 27 distinct categories of emotion bridged by continuous gradients. Proceedings of the National Academy of Sciences 114, E7900–E7909.

Craig, A., 2009. How do you feel-now? The anterior insula and human awareness. Nat Rev Neurosci. 10, 59–71. doi:10.1038/nrn2555.

Delorme, A., Makeig, S., 2004. Eeglab: an open source toolbox for analysis of single-trial eeg dynamics including independent component analysis. Journal of neuroscience methods 134, 9–21.

Delorme, A., Palmer, J., Onton, J., Oostenveld, R., Makeig, S., 2012. Independent eeg sources are dipolar. PloS one 7, e30135.

Delorme, A., Sejnowski, T., Makeig, S., 2007. Enhanced detection of artifacts in eeg data using higher-order statistics and independent component analysis. Neuroimage 34, 1443–1449.

Ekman, P., 1993. Facial expression and emotion. American psychologist 48, 384.

Horikawa, T., Cowen, A.S., Keltner, D., Kamitani, Y., 2020. The neural representation of visually evoked emotion is high-dimensional, categorical, and distributed across transmodal brain regions. iScience, 101060.

Hsu, S.H., Pion-Tonachini, L., Palmer, J., Miyakoshi, M., Makeig, S., Jung, T.P., 2018a. Modeling brain dynamic state changes with adaptive mixture independent component analysis. NeuroImage.

Hsu, S.H., Zi, Y., Wu, Y.C., Jackson, P.M., Jung, T.P., 2018b. Exploring mental state changes during hypnotherapy using adaptive mixture independent component analysis of eeg, in: 2018 IEEE Biomedical Circuits and Systems Conference (BioCAS), IEEE. pp. 1–4.

Jung, T.P., Makeig, S., Lee, T.W., McKeown, M.J., Brown, G., Bell, A.J., Sejnowski, T.J., 2000. Independent component analysis of biomedical signals, in: Proc. Int. Workshop on Independent Component Analysis and Signal Separation, Citeseer. pp. 633–644.

Kerestes, R., Ladouceur, C.D., Meda, S., Nathan, P.J., Blumberg, H.P., Maloney, K., Ruf, B., Saricicek, A., Pearlson, G.D., Bhagwagar, Z., Phillips, M.L., 2012. Abnormal prefrontal activity subserving attentional control of emotion in remitted depressed patients during a working memory task with emotional distracters. Psychological Medicine 42, 29–40. doi:10.1017/S0033291711001097.

Kim, M.K., Kim, M., Oh, E., Kim, S.P., 2013. A review on the computational methods for emotional state estimation from the human eeg. Computational and mathematical methods in medicine 2013.

Kober, H., Barrett, L.F., Joseph, J., Bliss-Moreau, E., Lindquist, K., Wager, T.D., 2008. Functional grouping and cortical-subcortical interactions in emotion: a meta-analysis of neuroimaging studies. Neuroimage 42, 998–1031.

Koch, S.B., Mars, R.B., Toni, I., Roelofs, K., 2018. Emotional control, reappraised. Neuroscience and Biobehavioral Reviews 95, 528–534. URL: https://doi.org/10.1016/j.neubiorev.2018.11.003, doi:10.1016/j.neubiorev.2018.11.003.

Koelstra, S., Muhl, C., Soleymani, M., Lee, J.S., Yazdani, A., Ebrahimi, T., Pun, T., Nijholt, A., Patras, I., 2011. Deap: A database for emotion analysis; using physiological signals. IEEE transactions on affective computing 3, 18–31.

Kothe, C.A., Makeig, S., Onton, J.A., 2013. Emotion recognition from eeg during self-paced emotional imagery, in: Proceedings of the 2013 Humaine Association Conference on Affective Computing and Intelligent Interaction, pp. 855–858.

Kothe, C.A.E., Jung, T.P., 2016. Artifact removal techniques with signal reconstruction. US Patent App. 14/895,440.

Lee, T.W., Lewicki, M., Sejnowski, T., 2000. Ica mixture models for unsupervised classification of non-gaussian classes and automatic context switching in blind signal separation. IEEE Transactions on Pattern Analysis and Machine Intelligence 22, 1078–1089. doi:10.1109/34.879789.

Leech, R., Sharp, D.J., 2014. The role of the posterior cingulate cortex in cognition and disease. Brain 137, 12–32. doi:10.1093/brain/awt162.

Liang, Z., Oba, S., Ishii, S., 2019. An unsupervised eeg decoding system for human emotion recognition. Neural Networks 116, 257–268.

Lin, Y.P., Wang, C.H., Jung, T.P., Wu, T.L., Jeng, S.K., Duann, J.R., Chen, J.H., 2010. Eeg-based emotion recognition in music listening. IEEE Transactions on Biomedical Engineering 57, 1798–1806.

Lindquist, K.A., Wager, T.D., Kober, H., Bliss-Moreau, E., Barrett, L.F., 2012. The brain basis of emotion: a meta-analytic review. The Behavioral and brain sciences 35, 121.

van der Maaten, L., Hinton, G., 2008. Visualizing data using t-SNE. Journal of Machine Learning Research 9, 2579–2605. URL: http://www.jmlr.org/papers/v9/vandermaaten08a.html.

Mauss, I.B., Robinson, M.D., 2009. Measures of emotion: A review. Cognition and emotion 23, 209–237.

Morawetz, C., Bode, S., Baudewig, J., Kirilina, E., Heekeren, H.R., 2016. Changes in Effective Connectivity Between Dorsal and Ventral Prefrontal Regions Moderate Emotion Regulation. Cerebral Cortex 26, 1923–1937. doi:10.1093/cercor/bhv005.

Morawetz, C., Bode, S., Derntl, B., Heekeren, H.R., 2017. The effect of strategies, goals and stimulus material on the neural mechanisms of emotion regulation: A meta-analysis of fMRI studies. Neuroscience and Biobehavioral Reviews 72, 111–128. URL: http://dx.doi.org/10.1016/j.neubiorev.2016.11.014, doi:10.1016/j.neubiorev.2016.11.014.

Motomura, K., Terasawa, Y., Natsume, A., Iijima, K., Chalise, L., Sugiura, J., Yamamoto, H., Koyama, K., Wakabayashi, T., Umeda, S., 2019. Anterior insular cortex stimulation and its effects on emotion recognition. Brain Structure and Function 224, 2167–2181. URL: https://doi.org/10.1007/s00429-019-01895-9, doi:10.1007/s00429-019-01895-9.

Mullen, T.R., Kothe, C.A., Chi, Y.M., Ojeda, A., Kerth, T., Makeig, S., Jung, T.P., Cauwenberghs, G., 2015. Real-time neuroimaging and cognitive monitoring using wearable dry eeg. IEEE Transactions on Biomedical Engineering 62, 2553–2567.

Nie, D., Wang, X.W., Shi, L.C., Lu, B.L., 2011. Eeg-based emotion recognition during watching movies, in: 2011 5th International IEEE/EMBS Conference on Neural Engineering, IEEE. pp. 667–670.

Onton, J.A., Makeig, S., 2009. High-frequency broadband modulation of electroencephalographic spectra. Frontiers in human neuroscience 3, 61.

Palmer, J.A., Makeig, S., Kreutz-Delgado, K., Rao, B.D., 2008. Newton method for the ica mixture model, in: Acoustics, Speech and Signal Processing, 2008. ICASSP 2008. IEEE International Conference on, IEEE. pp. 1805–1808.

Picard, R.W., 2000. Affective computing. MIT press.

Pion-Tonachini, L., Kreutz-Delgado, K., Makeig, S., 2019. Iclabel: An automated electroencephalographic independent component classifier, dataset, and website. NeuroImage 198, 181–197.

Ramnani, N., Owen, A.M., 2004. Anterior prefrontal cortex: Insights into function from anatomy and neuroimaging. Nature Reviews Neuroscience 5, 184–194. doi:10.1038/nrn1343.

Ran, S., Hsu, S.H., Jung, T.P., in press. Examining the relationship between eeg dynamics and emotion ratings during video watching using adaptive mixture independent component analysis. IEEE Conference on Systems, Man, and Cybernetics.

Ray, R.D., Zald, D.H., 2012. Anatomical insights into the interaction of emotion and cognition in the prefrontal cortex. Neuroscience and Biobehavioral Reviews 36, 479–501. doi:10.1016/j.neubiorev.2011.08.005.

Rolls, E.T., 2019. The cingulate cortex and limbic systems for emotion, action, and memory. Brain Structure and Function 224, 3001–3018. URL: https://doi.org/10.1007/s00429-019-01945-2, doi:10.1007/s00429-019-01945-2.

Safont, G., Salazar, A., Vergara, L., Gomez, E., Villanueva, V., 2017. Probabilistic distance for mixtures of independent component analyzers. IEEE Transactions on Neural Networks and Learning Systems, 1–13 doi:10.1109/TNNLS.2017.2663843.

Salazar, A., Vergara, L., Miralles, R., 2010. On including sequential dependence in ica mixture models. Signal Processing 90, 2314–2318. doi:10.1016/j.sigpro.2010.02.010.

Scherer, K.R., 2005. What are emotions? and how can they be measured? Social science information 44, 695–729.

Seghier, M.L., 2013. The angular gyrus: Multiple functions and multiple subdivisions. Neuroscientist 19, 43–61. doi:10.1177/1073858412440596.

Sequeira, H., Hot, P., Silvert, L., Delplanque, S., 2009. Electrical autonomic correlates of emotion. International journal of psychophysiology 71, 50–56.

Sivagnanam, S., Majumdar, A., Yoshimoto, K., Astakhov, V., Bandrowski, A.E., Martone, M.E., Carnevale, N.T., 2013. Introducing the neuroscience gateway. IWSG 993.

Viviani, R., 2014. Neural Correlates of Emotion Regulation in the Ventral Prefrontal Cortex and the Encoding of Subjective Value and Economic Utility. Frontiers in Psychiatry 5, 1–11. doi:10.3389/fpsyt.2014.00123.

Zhang, Y., Zhou, W., Wang, S., Zhou, Q., Wang, H., Zhang, B., Huang, J., Hong, B., Wang, X., 2019. The Roles of Subdivisions of Human Insula in Emotion Perception and Auditory Processing. Cerebral Cortex 29, 517–528. doi:10.1093/cercor/bhx334.

Zheng, W.L., Zhu, J.Y., Peng, Y., Lu, B.L., 2014. Eeg-based emotion classification using deep belief networks, in: 2014 IEEE International Conference on Multimedia and Expo (ICME), IEEE. pp. 1–6.

Zhuang, X., Rozgic, V., Crystal, M., 2014. Compact unsupervised eeg response representation for emotion recognition, in: IEEE-EMBS international conference on Biomedical and Health Informatics (BHI), IEEE. pp. 736–739.

