## Supplemental Materials for "Unsupervised Learning of Brain State Dynamics during Emotion Imagery using High-Density EEG"

### S1. Supplemental Methods

#### *S1.1. Adaptive Mixture Independent Component Analysis*

Comprehensive formulation and extensive evaluation of the AMICA algorithm can be found in Palmer et al. (2008) and Hsu et al. (2018) respectively. Conceptually, AMICA consists of three layers of mixing processes: AMICA learns a mixture of ICA models (Eq. 1); each model is a mixture of independent components (IC) or sources, and each IC has a probability density function parameterized as a mixture of generalized Gaussians (Palmer et al., 2008).

The first layer assumes that data $\boldsymbol{x}$ ($N$-channels $\times$ $T$-samples) are nonstationary, so that different models may best characterize the data at different times, i.e., $\boldsymbol{x}(t) = \boldsymbol{x}_h(t)$ where $h$ is the model index. In the second layer, a standard ICA model is employed to model the data $\boldsymbol{x}$ as an instantaneous linear mixture $\boldsymbol{A}$ ($N$-channels $\times$ $N$-sources) of statistically independent components $\boldsymbol{s}$ ($N$-sources $\times$ $T$-samples), i.e., $\boldsymbol{x} = \boldsymbol{A}\boldsymbol{s}$. The first two layers constitute the ICA mixture model:

$$\boldsymbol{x}(t) = \boldsymbol{x}_h(t) = \boldsymbol{A}_h \boldsymbol{s}_h(t) + \boldsymbol{b}_h, \quad h = 1, \ldots, H \tag{1}$$

where $h = h(t)$ and $\boldsymbol{A}_h$ is the dominant or active model at time $t$ with source activities $\boldsymbol{s}_h(t)$ and bias $\boldsymbol{b}_h$. Assuming $\boldsymbol{x}(t)$ are temporally independent, the likelihood of data given the ICA mixture model can be written as:

$$p(\boldsymbol{X}|\Theta) = \prod_{t=1}^{T} \sum_{h=1}^{H} p(\boldsymbol{x}(t)|C_h, \theta_h) \cdot p(C_h) \tag{2}$$

where $\Theta = \{\theta_1, \ldots, \theta_H\}$ contains the parameters of ICA models and $p(C_h)$ is the probability of the $h$-model being active that satisfies $\sum_{h=1}^{H} p(C_h) = 1$.

As an unsupervised approach with generative models, the $\Theta$ parameters learned by AMICA provide rich information about the underlying data clusters and their temporal dynamics. Specifically, the activation of each ICA model $h(t)$ can be represented as the data likelihood given the estimated parameters of each model $\theta_h$, i.e., $L_{h(t)} = p\big(\boldsymbol{x}(t)\big|\theta_{h(t)}\big)$. Therefore, the probability of activation of each ICA model at time $t$ can be calculated by normalizing $L_{h(t)}$ across all models and is referred to as "ICA model probability" that indicates the goodness-of-fit of the ICA model to the data samples (Hsu et al., 2018).

The expectation-maximization (EM) algorithm is employed to estimate the parameters $\hat{\Theta}$ that maximize the data likelihood function in Eq. 2. In the M-step, AMICA uses the Newton approach based on the Hessian (matrix of second-order derivatives) to achieve faster convergence.

Multi-model AMICA decomposition was applied to each EEG dataset with the following parameters: number of models $= 20,$ number of generalized gaussians mixture $= 1$, and max learning iterations $= 2000$. Data samples with low probabilities of model fit were rejected ($numreg = 5, rejstart = 2, rejint = 5$) from being used for learning AMICA parameters to alleviate the effects of transient artifacts such as electrode pops and discontinuities. A sphering transformation was applied prior to AMICA decomposition ($do\_pca = 1$). The AMICA jobs were run on Comet Supercomputer, San Diego Supercomputer, using one computing node with 24 threads ($max\_threads = 24$) under the support of Neuroscience Gateway (NSG) (Sivagnanam et al., 2013).

An efficient implementation of AMICA with parallel computing capability by Palmer et al. (2008) was used in this study. The code for the implementation is available at `https://github.com/japalmer29/amica` and as an open-source plug-in for EEGLAB (Delorme and Makeig, 2004). For detailed instructions and tutorials of running AMICA, please visit `https://sccn.ucsd.edu/wiki/AMICA`

The output parameters of AMICA are 20 ICA models (i.e., mixing matrices $\boldsymbol{A}_h$ in Eq. 1) and their model-probability time courses (i.e., normalized data likelihood). Since brain activity is non-stationary throughout the emotion imagery experiment, we expect the probability time course of each model would fluctuate, reflecting changes of EEG patterns captured by different models.

### S2. Supplemental Table

Table S1: Distribution of types of AMICA models for each subject.

| Subject ID \ Type | I | II | III | IV |
|---|---|---|---|---|
| 1 | 5 | 0 | 0 | 3 |
| 2 | 4 | 6 | 1 | 1 |
| 4 | 2 | 0 | 8 | 1 |
| 5 | 2 | 2 | 2 | 1 |
| 6 | 4 | 6 | 3 | 1 |
| 7 | 2 | 4 | 3 | 1 |
| 8 | 3 | 4 | 3 | 1 |
| 9 | 6 | 6 | 0 | 1 |
| 10 | 3 | 5 | 3 | 0 |
| 11 | 6 | 4 | 1 | 1 |
| 12 | 3 | 0 | 2 | 3 |
| 13 | 0 | 2 | 0 | 2 |
| 14 | 6 | 1 | 1 | 0 |
| 15 | 4 | 2 | 4 | 1 |
| 16 | 2 | 3 | 1 | 2 |
| 17 | 4 | 7 | 2 | 0 |
| 18 | 6 | 2 | 2 | 1 |
| 19 | 1 | 1 | 2 | 4 |
| 20 | 5 | 4 | 2 | 1 |
| 21 | 2 | 4 | 2 | 3 |
| 23 | 4 | 5 | 3 | 2 |
| 24 | 2 | 3 | 0 | 2 |
| 25 | 1 | 6 | 4 | 1 |
| 27 | 5 | 3 | 4 | 0 |
| 28 | 5 | 5 | 1 | 0 |
| 29 | 2 | 7 | 1 | 0 |
| 31 | 4 | 1 | 8 | 0 |
| 33 | 3 | 0 | 3 | 3 |
| 35 | 3 | 1 | 0 | 4 |

### S3. Supplemental Figures

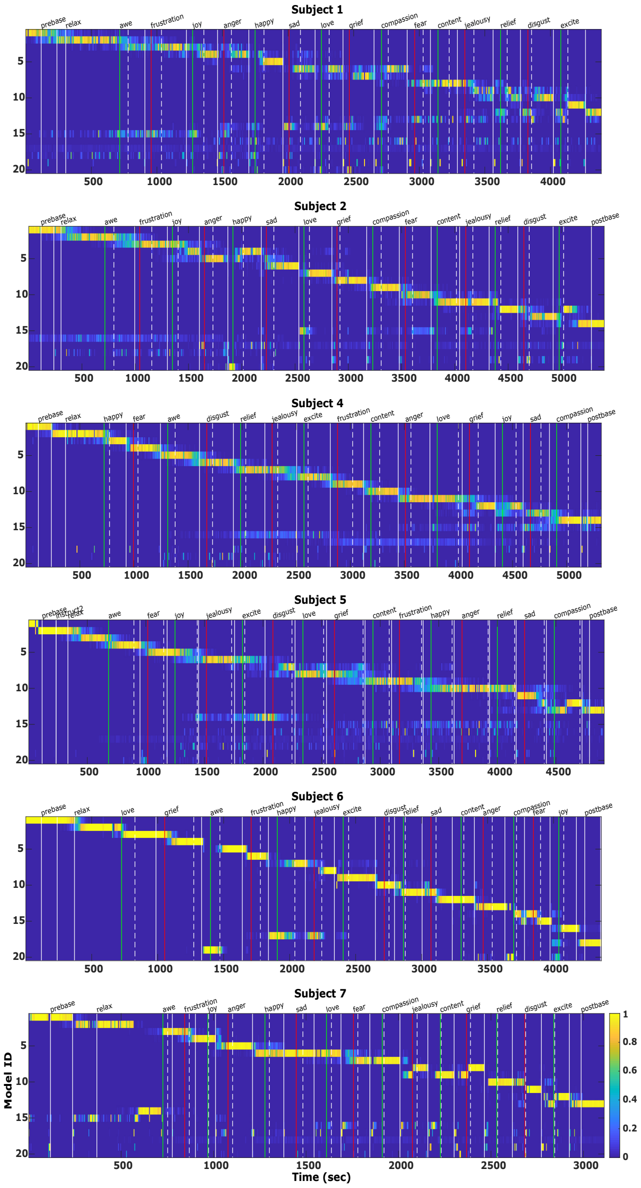


Figure S1: Probability time courses of 20 models learned by AMICA for all of the 29 subjects (Fig. S1 to Fig. S5). The models were sorted based on their active time in the experiment. The vertical color lines indicate the start of each emotion imagery trial with positive (green) and negative (red) emotions. The dashed and solid white lines indicate the button presses when the subject felt the emotion and when the subject exited the trial.

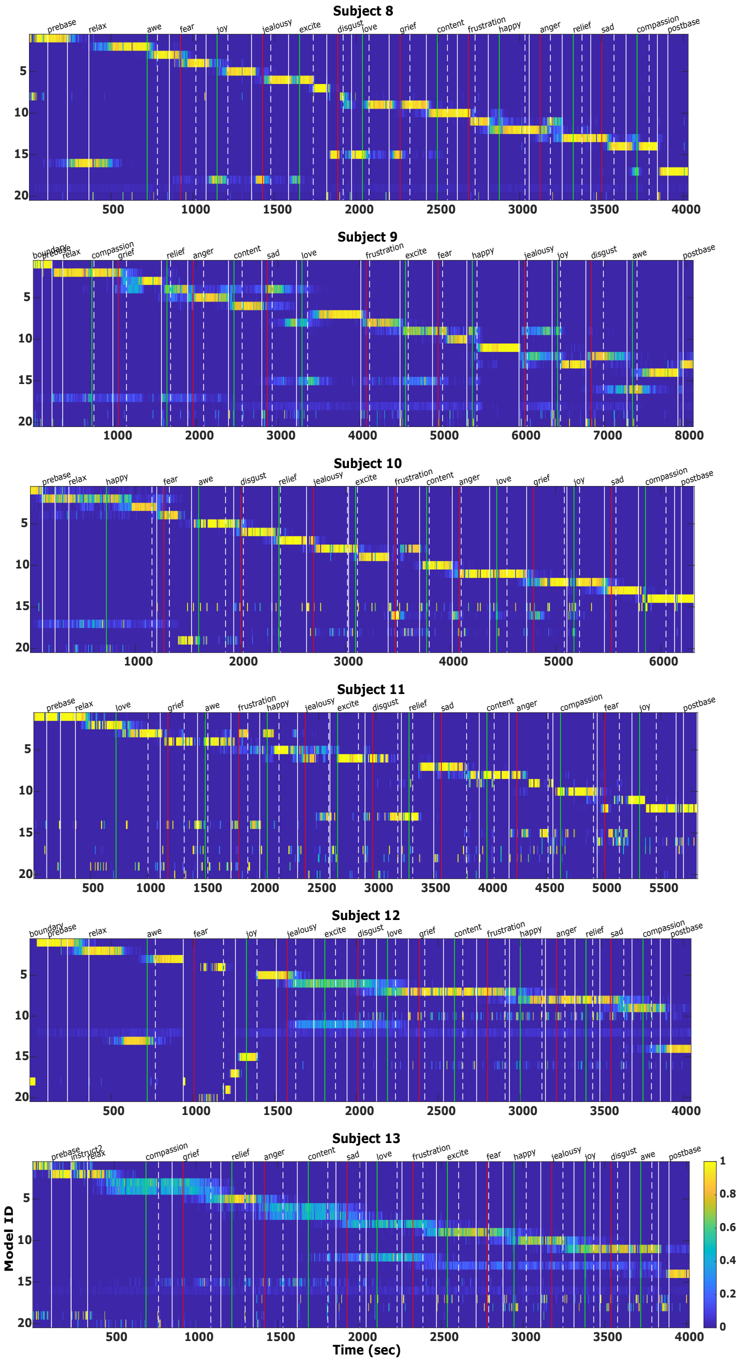

Figure S2: Continued from Fig. S1

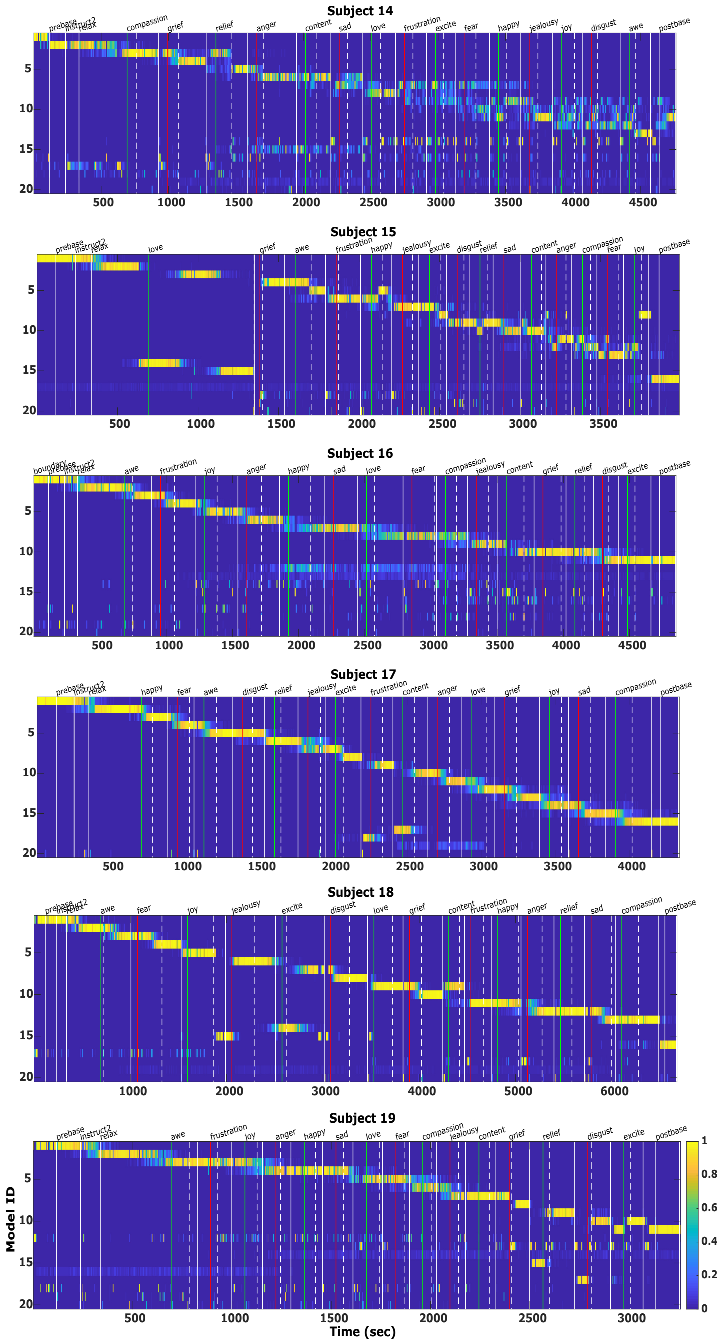

Figure S3: Continued from Fig. S2

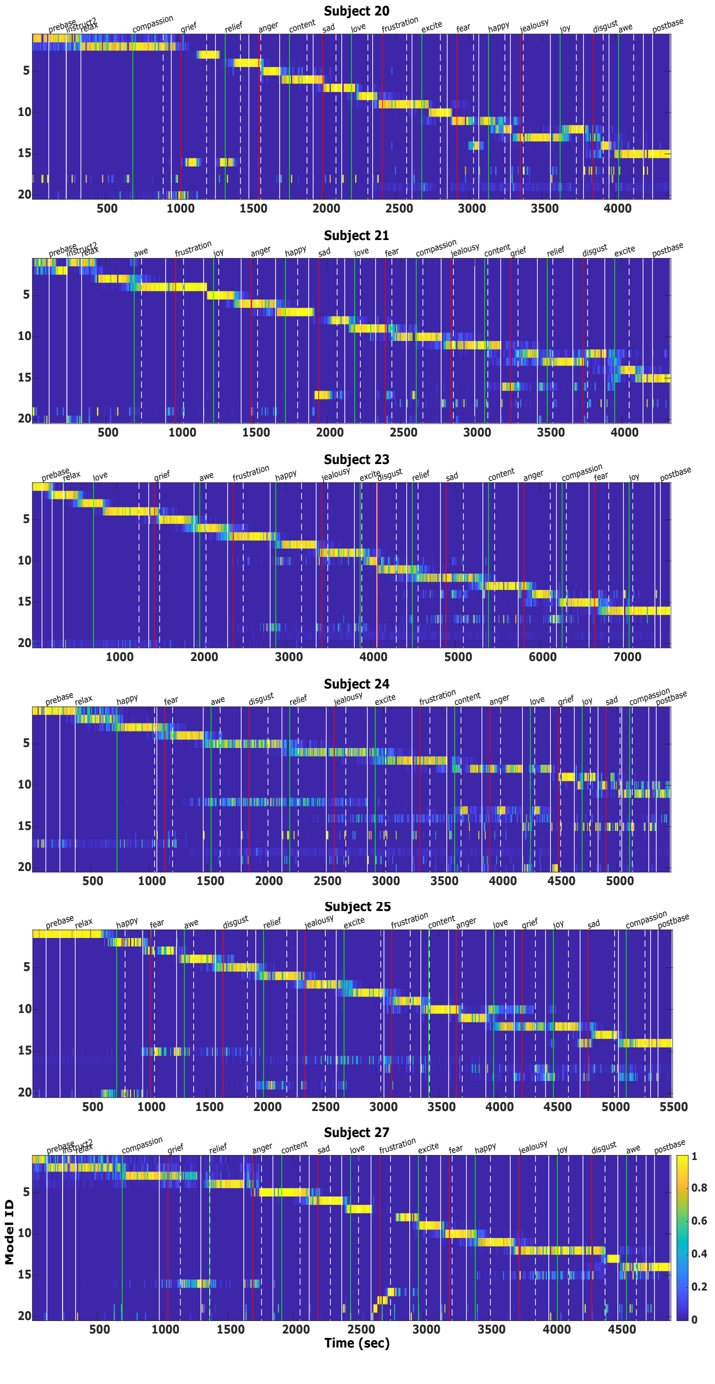


Figure S4: Continued from Fig. S3

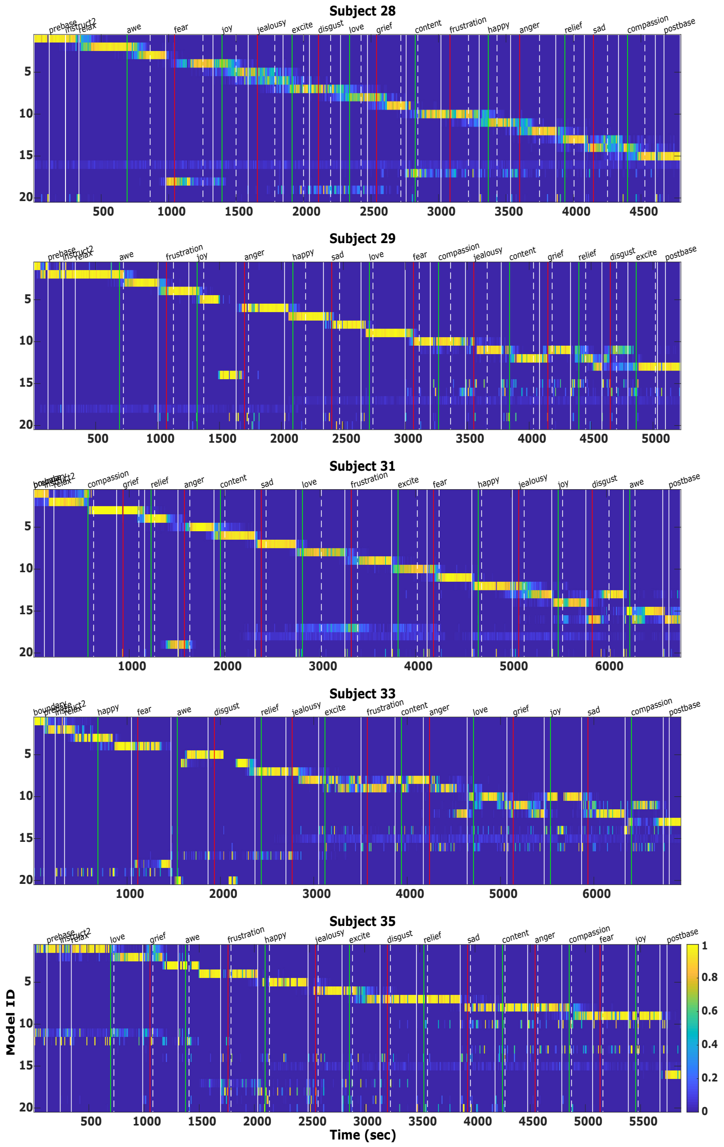


Figure S5: Continued from Fig. S4

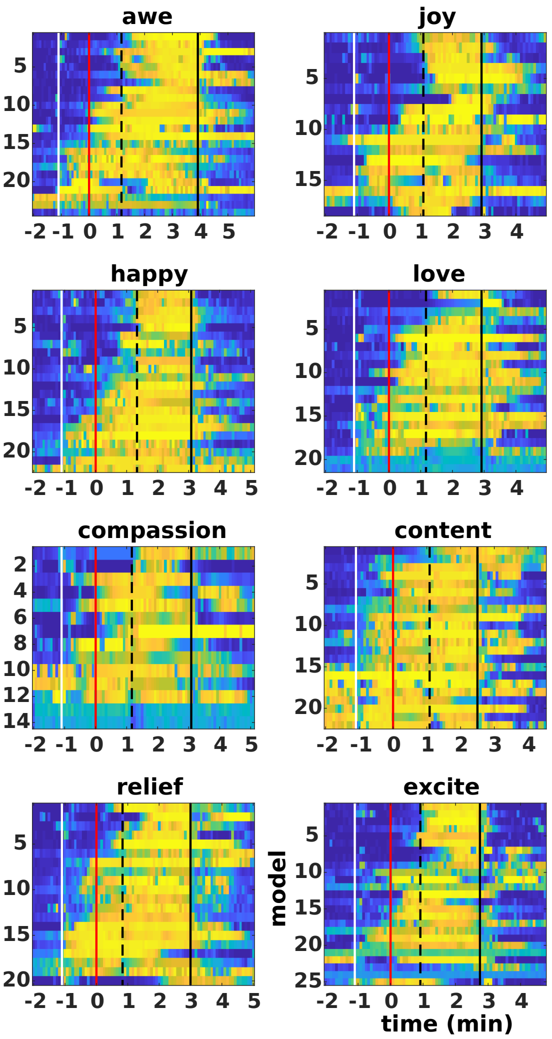


Figure S6: The model-probability time-series of AMICA models across subjects in the same cluster activated during each of the 15 emotions, time-locked to the beginning of the trials and smoothed with a 5-sec non-overlapping window. The vertical lines mark the exit from the previous emotion trial and start of the resting period (white) and the onset of the audio instruction (red). The first button presses (dashed line) and the end of the current emotion trials (dash-dot line) were time-warped to the median reaction time of all subjects whose model contributed to the cluster. The models were sorted according to their types defined by the timing of being active.

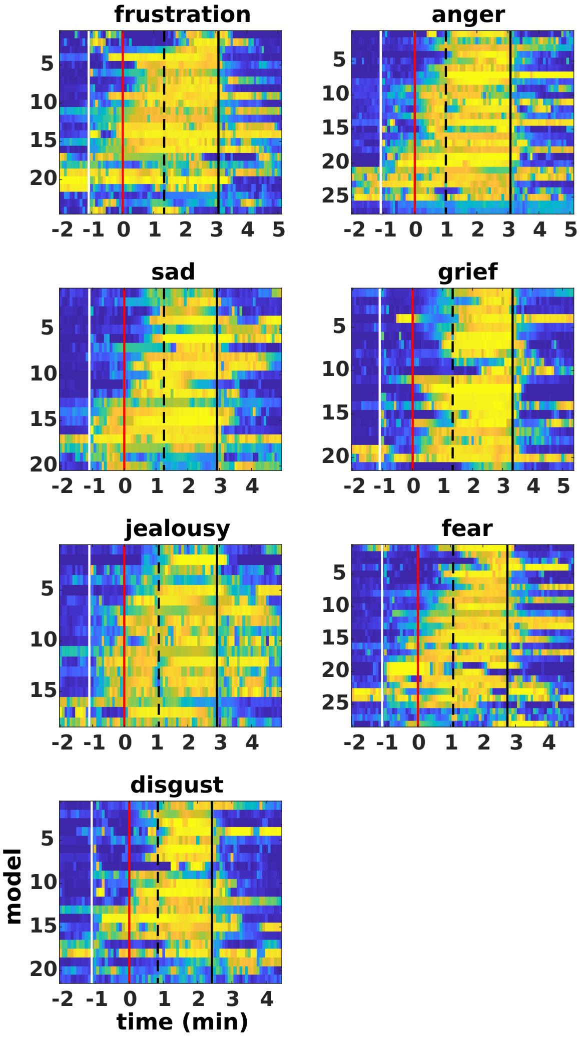


Figure S7: Continued from Fig. S6

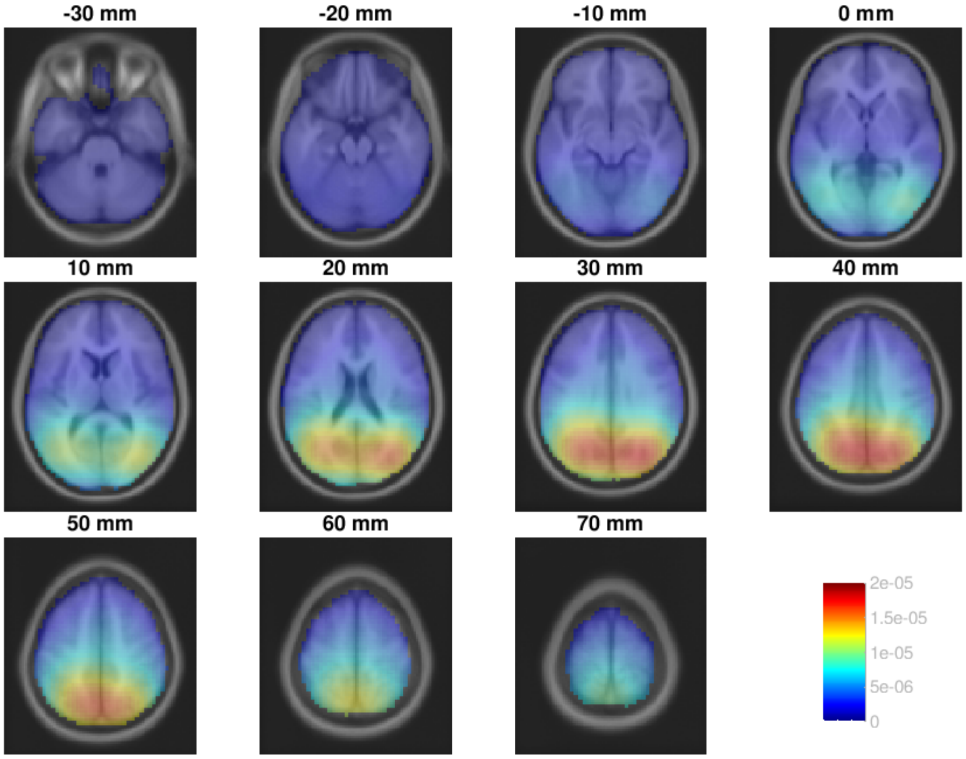


Figure S8: The dipole density of dipolar brain ICs of all AMICA models activated during baseline periods (i.e. pre- and post-session baseline and relaxation), superimposed on the axial slices of a template MR image.

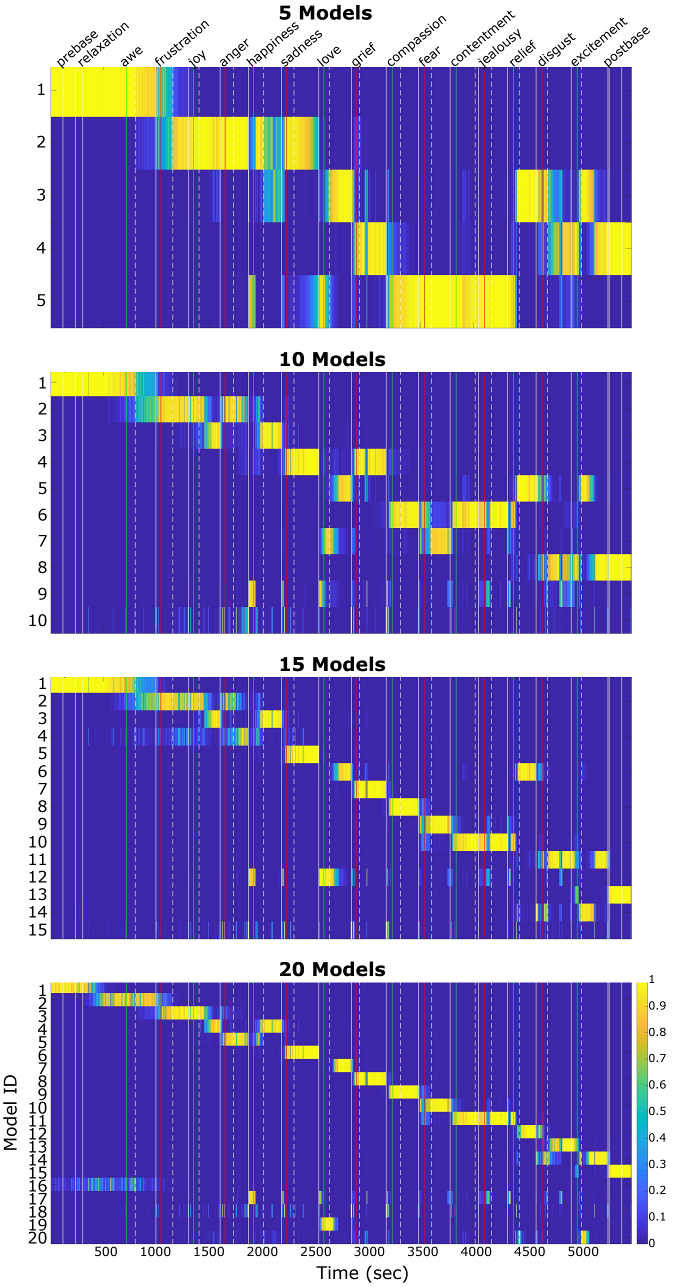


Figure S9: Probability time courses of 5, 10, 15, and 20 models learned by AMICA for a representative subject. The models were sorted based on their active time in the experiment. The vertical color lines indicate the start of each emotion imagery trial with positive (green) and negative (red) emotions. The dashed and solid white lines indicate the button presses when the subject felt the emotion and when the subject exited the trial.

### References


Delorme, A., Makeig, S., 2004. Eeglab: an open source toolbox for analysis of single-trial eeg dynamics including independent component analysis. Journal of neuroscience methods 134, 9–21.

Hsu, S.H., Pion-Tonachini, L., Palmer, J., Miyakoshi, M., Makeig, S., Jung, T.P., 2018. Modeling brain dynamic state changes with adaptive mixture independent component analysis. NeuroImage .

Palmer, J.A., Makeig, S., Kreutz-Delgado, K., Rao, B.D., 2008. Newton method for the ica mixture model, in: Acoustics, Speech and Signal Processing, 2008. ICASSP 2008. IEEE International Conference on, IEEE. pp. 1805–1808.

Sivagnanam, S., Majumdar, A., Yoshimoto, K., Astakhov, V., Bandrowski, A.E., Martone, M.E., Carnevale, N.T., 2013. Introducing the neuroscience gateway. IWSG 993.
